# CELLNET technology: Spatially organized, functional 3D networks at single cell resolution

**DOI:** 10.1101/2024.07.12.603216

**Authors:** Arun Poudel, Puskal Kunwar, Ujjwal Aryal, Anna-Blessing Merife, Pranav Soman

## Abstract

Cells possess the remarkable ability to generate tissue-specific 3D interconnected networks and respond to a wide range of stimuli. Understanding the link between the spatial arrangement of individual cells and their networks’ emergent properties is necessary for the discovery of both fundamental biology as well as applied therapeutics. However, current methods spanning from lithography to 3D photo-patterning to acoustofluidic devices are unable to generate interconnected and organized single cell 3D networks within native extracellular matrix (ECM). To address this challenge, we report a novel technology coined as CELLNET. This involves the generation of crosslinked collagen within multi-chambered microfluidic devices followed by femtosecond laser ablation of 3D microchannel networks and cell seeding. Using model cells, we show that cell migrate within ablated networks within hours, self-organize and form viable, interconnected, 3D networks in custom architectures such as square grid, concentric circle, parallel lines, and spiral patterns. Heterotypic CELLNETs can also be generated by seeding multiple cell types in side-chambers of the devices. The functionality of cell networks can be studied by monitoring the real-time calcium signaling response of individual cells and signal propagation within CELLNETs when subjected to flow stimulus alone or a sequential combination of flow and biochemical stimuli. Furthermore, user-defined disrupted CELLNETs can be generated by lethally injuring target cells within the 3D network and analyzing the changes in their signaling dynamics. As compared to the current self-assembly based methods that exhibit high variability and poor reproducibility, CELLNETs can generate organized 3D single-cell networks and their real-time signaling responses to a range of stimuli can be accurately captured using simple cell seeding and easy-to-handle microfluidic devices. CELLNET, a new technology agnostic of cell types, ECM formulations, 3D cell-connectivity designs, or location and timing of network disruptions, could pave the way to address a range of fundamental and applied bioscience applications.

**Teaser:** New technology to generate 3D single cell interconnected and disrupted networks within natural extracellular matrix in custom configurations.

## Introduction

Recreating the 3D spatial organization of single-cell networks will help elucidate underlying mechanisms related to their emergent functional properties exhibited at the tissue level. Although technological advances have led to an array of methods to arrange cells in 2D and 3D to mimic tissue specific architectures, single-cell resolution with control over cell-to-cell connectivity remains challenging.^1–2^ A common strategy is to pattern 2D adhesive micropatterns using conventional lithography methods followed by cell seeding to generate interconnected single cell networks. Other methods such as laser-assisted bioprinting has also been used to directly deposit single cells and generate 2D cellular arrays, however, arranging single cells in 3D has proved difficult.^3–4^ Methods based on 3D photo-patterning of adhesive peptides within specialized semi-synthetic hydrogels have emerged,^5^ but they have failed to realize 3D single cell networks. At present, only methods based on multi-photon absorption (MPA) can pattern features at single-cell resolutions^6–15^ although limitations related to specialized photosensitive materials, water soluble low-toxicity photoinitiators, cell viability during laser scanning in the presence of cells, and low scalability, have limited its use in the field. Thus, the field continues to rely on self-assembly based methods, that involve mixing relevant cells in natural ECM like collagen or fibrin. This however results in randomly organized 3D cell networks without precise control over network density, connectivity, and architecture, making systematic mechanistic study on networks’ properties challenging. Bioprinting methods can provide spatial control over cell placements within 3D ECMs,^16–20^ however single cell resolution is not possible.^21^ Microfluidic devices integrated with acoustic,^22–24^ dielectric^25–26^, and magnetic field stimulation^26–27^ have also been used to directly manipulate cells in a contactless manner. However, these methods cannot achieve single-cell resolution or user-defined multi-layer 3D patterns, and its control over intercellular connectivity remains poor. In summary, current methods are unable to generate tissue-specific, 3D, single-cell networks within natural, unmodified extracellular matrix, like collagen. To address this challenge, we report a new technology, CELLNET, to generate 3D, single cell, functional networks in custom architectures within multi-chambered chips using laser ablation of 3D microchannel templates within collagen matrix. **(Fig.1)** First, Digital Light Projection (DLP) was used to rapidly design and print master molds which were used to generate custom three-chambered PDMS devices. Second, ECM of interest (type I collagen) was perfused into central chambers and thermally crosslinked to generate a barrier between chambers 1 and 3. Third, Two Photon Ablation (TPA) was used to ablate 3D microchannel network within collagen. Model cells, seeded within the device, self-assemble within ablated network, and generate an interconnected, 3D, functional circuits coined as CELLNETs. We show that CELLNETs are compatible with standard imaging methods (brightfield, immunostaining, time-lapse microscopy), co-culturing cells and in situ manipulation such as application of fluid flow and/or biochemical stimuli, or injury to target cells to generate user-defined disrupted networks. CELLNET, generated using tissue-specific cells, ECM, and single-cellular spatial organization, can be potentially used as a deterministic model to study how short-term signaling results in durable functional or yet unknown emergent properties.

**Fig. 1.**
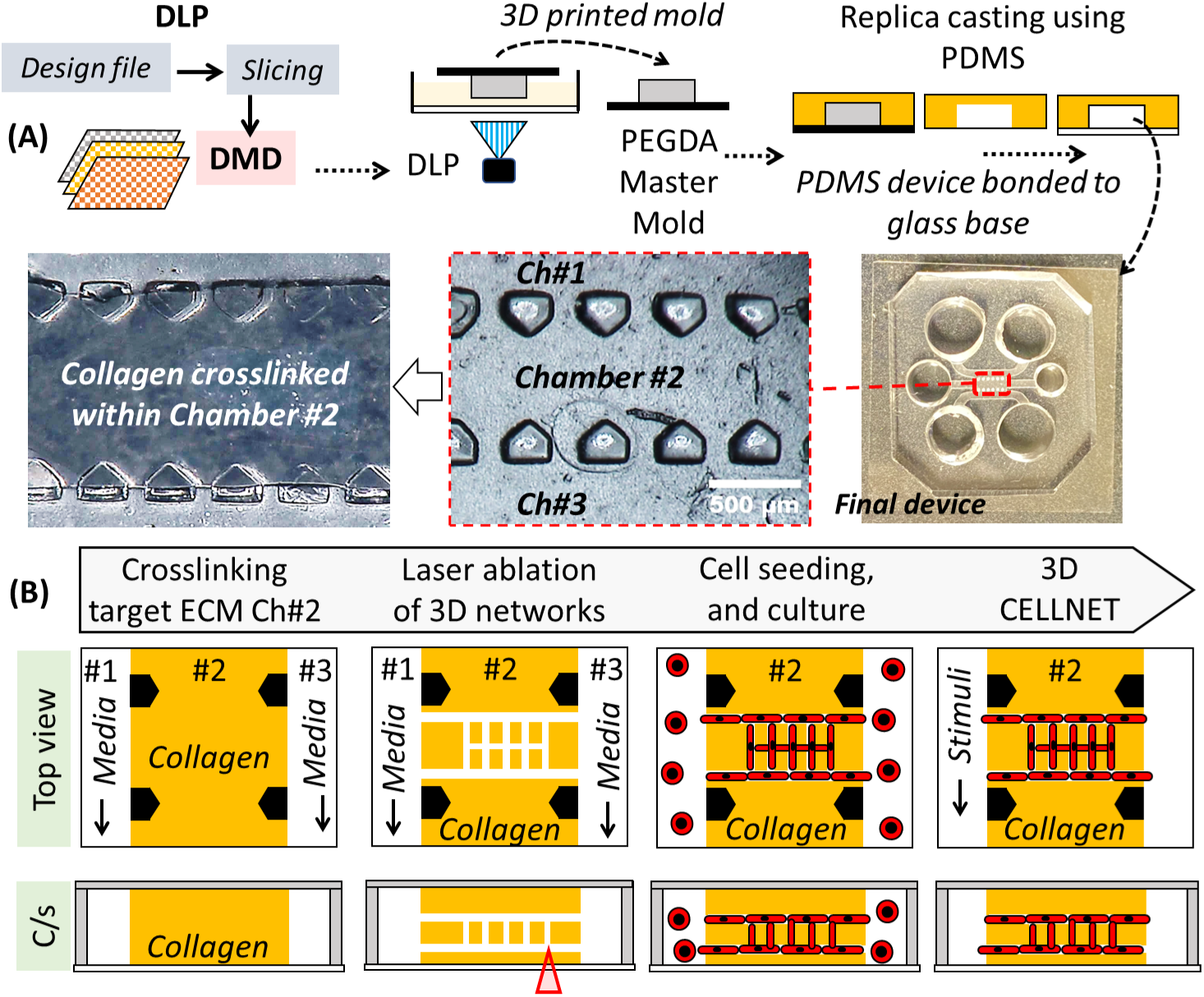
(A) Schematic showing 3D printing of Master molds using Digital Light Projection (DLP) followed by replica casting to generate three-chambered microfluidic devices using PDMS (picture). Type I collagen solution is perfused and crosslinked within chamber #2. (B) Process flow with top and cross-sectional views to generate a two-layer interconnected cellular networks or CELLNETs within Type I Collagen in the central chamber of the microfluidic device.

## Results

### Design and development of microfluidic devices using DLP-printed Master Molds

We used DLP to print master molds which were used to replica cast final PDMS microfluidic devices **(Fig.1A, SI-1A)**. Details related to the DLP printing setup, pre-polymer formulation, printing conditions, glass modification protocols, are reported in the Methods and SI sections. Briefly, a CAD model of a reverse master mold of the intended microfluidic device was designed in SolidWorks; this consist of positive features that would be replicated into three microfluidic chambers with negative spaces, each with inlet and outlet ports. **(SI-1B)**. CAD model was then sliced to generate a series of virtual masks which were converted to corresponding light patterns by the Digital Micromirror Device (DMD). Upon irradiation onto liquid PEGDA (250MW) photo-polymer solution in a layer-by-layer manner, final master mold was realized. Glass surface was modified to prevent delamination of PEGDA mold during the printing process. Molds were printed at the step size of 20µm with 0.8s exposure time per layer with constant light intensity of 6mWcm^−2^. Post-printing, molds were cleaned with 100% ethanol to remove any uncured resin. Molds were cured for an additional 20s under UV light to ensure strong crosslinking between the mold and the bottom glass slide to prevent any delamination during the PDMS casting steps. PDMS prepolymer solution was casted onto the molds followed by thermally curing at low and high temperature cycles; this ensured high repeatability, since without the low temperature curing step, molds show tendency to crack and warp. Prior to casting, PEGDA molds were submersed in 100% ethanol for an hour followed by exposure to ambient light for 24 hours to generate defect-free PDMS devices. Without these post-processing steps, PDMS casting led to many defects possibly due to leaching of free radicals from the interior of the mold causing inhibition of PDMS crosslinking at the mold-liquid PDMS interface.^28^**(SI-1C)** Polymerized PDMS was carefully peeled off, trimmed to suitable size, and 6 inlet-outlet holes were punched (2 holes per chamber) before irreversibly bonding to a glass coverslip (0.15 mm thick) to generate final PDMS devices. **(Fig. 1A)** The final devices consist of three-chambers; a central chamber (Ch#2; 0.85mm wide, to house crosslinked collagen) flanked on either side by two chambers (Ch#1 and Ch#2; 1mm wide, for media exchanges), separated by an array of microposts (width = 200μm) with a spacing of 200μm. The height for all the chambers were 240µm.

### Characterization of laser ablated 3D microchannel networks within crosslinked collagen

In this work, we choose type I collagen as model ECM due to its abundance in *in vivo* tissues, and wide use in the field to generate 3D cell culture models. Before pipetting collagen solution within PDMS devices, they were surface modified to prevent delamination of crosslinked collagen from the channel surfaces during active cultures. First (3-aminopropyl)triethoxy silane (APTES) was used to silanize the glass/PDMS surfaces to generate self-assembled monolayer (SAM) with reactive functional groups (-NH2) followed by glutaraldehyde (GA) to form reactive functional groups (-COOH).^29^ Without these modification steps, the crosslinked collagen showed a tendency to detach from the PDMS roof and glass bottom surfaces. **(SI-1D)** Then, type I collagen solution (4mg/ml) was crosslinked within the central chambers of UV sterilized devices. **(Fig.1B)** After 24 hours, a focused femtosecond laser was used to ablate 3D microchannels architectures (800nm, 1.2W; ablation setup is shown in **SI-2A**). To reliably generate 3D channels of defined lumen sizes, laser ablation was performed at varying power and depths inside collagen (50 - 200µm) in both lateral (XY) **(Fig.2A, SI-2B, SI-Video 1)** and vertical (Z) **(Fig.2D, SI-2C, SI-Video 2)** directions. Reflectance microscopy images show top and cross-sectional views of ablated channels embedded in collagen matrix. Laser dosage below 2 x 10^5^ Jcm^-2^ results in ablated microchannels with lumen size of 0.5-1µm, while above this threshold, laser irradiation results in cavitation and formation of larger sized lumens (8-10µm). **(Fig.2B)** During laser scanning above this threshold, the radial expansion of the bubble generates a shockwave which locally breaks down the collagen network and leaves behind a hollow lumen in its wake. **(SI-3A,B)** We also noticed that gas bubbles were elongated in the scanning direction and often remain in the channel from 10s of seconds to 1-2 hours. We found that if ablation was carried out immediately after collagen crosslinking, scanning induced bubble collapses and the channels close, possibly due to partially crosslinked collagen. Thus, 24hrs were given for completion of collagen crosslinking before ablation was performed; this ensured repeatable and stable channel formation without collapse. **(SI-3C)** As expected, for constant laser dosage, lumen size decreased with increased depth in collagen. **(Fig.2C)** Laser processing plots (shown in **Fig.2B,D**) were used as a design guide to repeatably generate 3D microchannel networks with a lumen size of ∼8µm within collagen; this lumen size was found to be ideal for single cell migration. Lumen size >8µm led to migration of multiple cells within the channels while lumen size <8µm decreased single cell migration and resulted in network occupancy below 50%, as discussed in later sections. For this work, we used 4mg/ml collagen concentration with a laser power of 1.4 W and 40x water objective (0.8NA) to generate various configurations of CELLNETs. Fluorescent beads (∼0.5μm) were used to verify perfusion within 3D ablated channel network (lumen size∼8µm) within the central chamber (#2) of a microfluidic device. **(Fig.2E-G).**

**Fig. 2.**
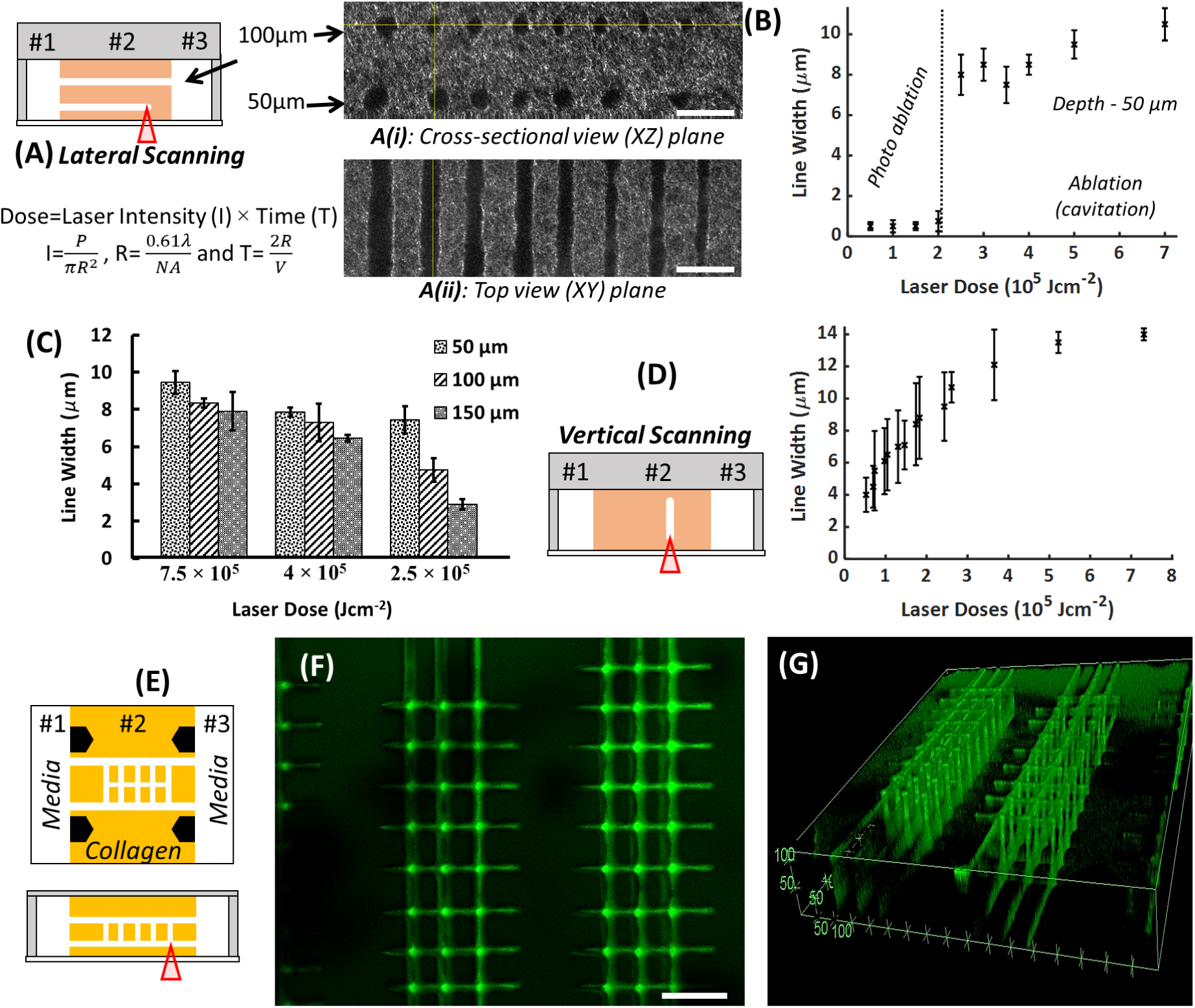
Characterization of microchannels ablated within collagen. (A) Schematic showing lateral scanning (XY) with varying dosage, and reflectance confocal images of ablated channels. Scale bar: 25μm (B) Plots showing the relationships between ablated line width and laser dosage at a depth of 50µm, while (C) shows the effect of varying depth on ablated line width. (D) Schematic and plot showing vertical ablation of microchannels and relationship between ablated line width (lumen size) and laser dosage. (E) Schematic of a 3D ablated grid with lumen size of ∼8µm in XY (lateral) and Z (vertical) directions within crosslinked collagen in chamber #2 of a three-chambered microfluidic device. (F-G). Top view and isometric confocal images after perfusion of a solution of fluorescent microbead (ϕ=0.5µm) into the ablated channels. Scale bar: 100μm.

### Generation of viable and interconnected single-cell 3D networks within microfluidic devices

We used fibroblast-like model 10T1/2 cells to demonstrate the formation of CELLNETs. First, cell solution (1M/ml) was pipetted in one of the side chambers (#1 or #3) of the microfluidic device, and brightfield microscopy was used to monitor cell migration within the ablated channels. We observed migration as early as 3 hours post-seeding. Between Day 1-2, cells migrated inside the collagen (500-1000μm from the interface between Ch#2 and Ch#1/3), and assemble into a 3D, interconnected single cell network within chamber #2 of the microfluidic device. **(Fig.3A, SI-4A)** On Day 2, extra cells were flushed out using 0.25% trypsin treatment for 5 minutes followed by pipetting fresh media in the side chambers; this step was performed to prevent aggregation of cells in Ch#1 or Ch#3 that could block diffusion of reporter dyes or antibodies and generate unwanted fluorescence during the characterization of cell networks. We observe that post-trypsin wash, cell processes within the microchannel network retract for a few hours before recovering back to their normal spread morphology after 12 hours. **(SI-4B)**

**Fig. 3.**
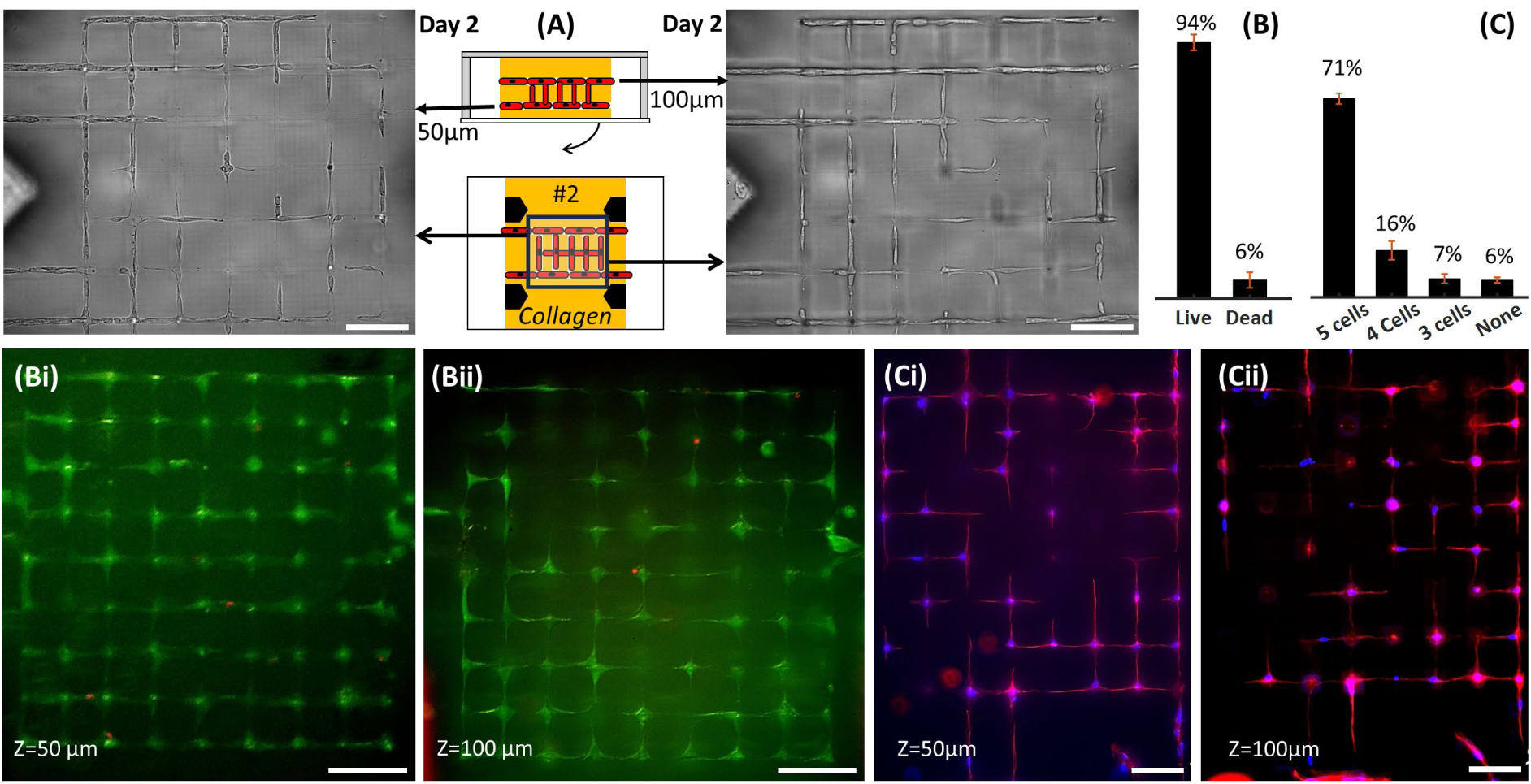
(A) Schematic and representative brightfield images from two z-planes (located at 50µm and 100µm from the bottom glass surface inside collagen matrix) showing seeding of model cells (10T1/2s) in chamber #1 or #3, and their migration within ablated networks to form viable, interconnected, 3D single-cell networks or CELLNETs. (Bi, ii) Viability of CELLNETs on Day 7 taken from both z-planes of the 3D network (C) Connectivity plot (Day 7, Ci, ii) Morphology of cells (actin=red; nucleus=blue).

To evaluate viability and morphology of cells within CELLNETs, a 3D microchannel network (lumen=8µm) with a two-layer connected grid architecture was used. The upper- and lower-layer, located 50µm and 100µm respectively from the glass bottom substrate, connect with each other. Viability of cells, calculated from cells from both layers, was found to be 93.23 ± 2.87% **(Fig.3B, i-ii)**. Fluorescence image of cells fixed on DAY 7 shows morphology (actin/nucleus) of cells assembled into pre-templated square-grid microchannel networks that provide defined control over cell-to-cell connectivity within collagen. **(Fig.3Ci-ii)** For instance, in the template used here, for a target cell within the CELLNET, a maximum of 5 cell-cell connections can be generated: 4 in-plane connections (in either the upper or lower layers) and one-out of plane (between the two layers). **(Fig.3C)** Results show that 70.56 ± 1.92 % cells were connected to 5 cells, while 16.67 ± 3.33 % were connected to 4 cells, 6.67 ± 1.67 % were connected to 3 cells, while 5 ± 0.96 % were not connected to any cells. This shows that around 95% cells within CELLNETs are connected to at least three neighboring cells. **SI-5** shows composite brightfield and fluorescence images, and a reconstructed 3D image.

To demonstrate that CELLNETs of complex topology can be generated, we designed microchannel templates such as parallel lines, spider webs, out-of-plane spirals, and double helixes. **(Fig.4)** Results show that cells migrate and self-organized into single cell 3D networks in custom orientations.

**Fig. 4.**
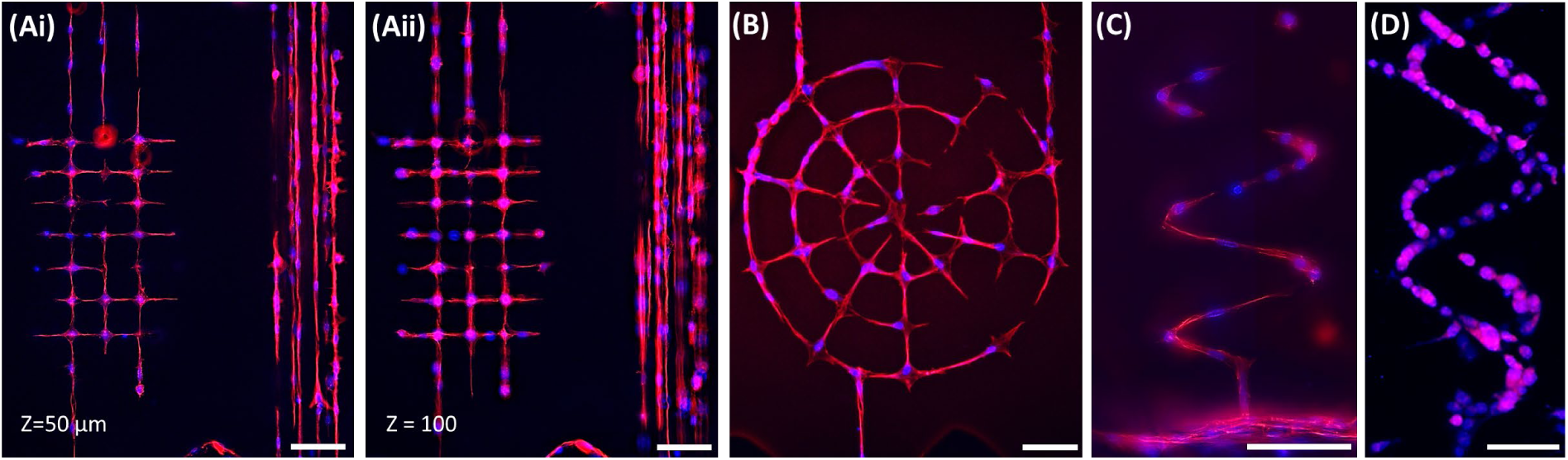
(A) Representative images showing single cells organized within user-defined square grid and parallel line architectures in two different z-planes within collagen matrix. Representative image of cell morphology organized in various patterns (B) concentric rings, (C) out-of-plane spiral and (D) double-helix. Scale bar: 100µm. (Actin: Red; Nucleus: Blue)

To demonstrate that CELLNETs can work with many cell types, we generated 3D square-grid microchannel templates within collagen, and seeded fibroblasts-like 10T1/2s, MLO-Y4 osteocytes, and Saos-2 osteoblast-like cells. Top, side and 3D reconstructed views of CELLNETs show formation of CELLNETs with in-plane and out-of-plane connections. **(Fig.5A-C)** Figure 5Ai highlights a cross-sectional section ∼75µm from the bottom glass substrate – a plane between the upper and lower microchannel grids showing ∼80% of out-of-plane actin-labelled cellular connections that span ∼50µm distance in the vertical direction. High resolution images show that some regions within the CELLNETs house more than one cells; nodes where 4 microchannels meet tend to slightly larger than the channels, thus have a higher likelihood of housing more than one cells. We found that cell connectivity is a function of lumen size and cell type and need to be optimized to get a single cell connectivity close to 100%. **(SI-Fig.6 Aiii and Bii)** For instance, for MLO-Y4 CELLNETs, an ablated channel size of ∼8µm ensuring single cell occupancy with ∼85% cell-to-cell connectivity while a lumen size of ∼12µm increases connectivity to ∼97% at the cost of having more than one cells in the larger ablated channels. Furthermore, we tested the capability of generating CELLNETs using two different cell types by seeding cells tagged with different fluorescent dyes in either side of the central chamber (Ch#2) **(SI-Fig.7)**. A 3D square-grid microchannel network was ablated within collagen. One study involved fluorescently tagging of the same cell type (MLO-Y4) with green and red dye, while the other study tagged MLO-Y4 with green dye and pre-osteoblasts (MC3T3) with red dye. In both cases, cells migrate towards each other within the 3D ablated microchannel network and form heterotypic CELLNETs.

**Fig. 5.**
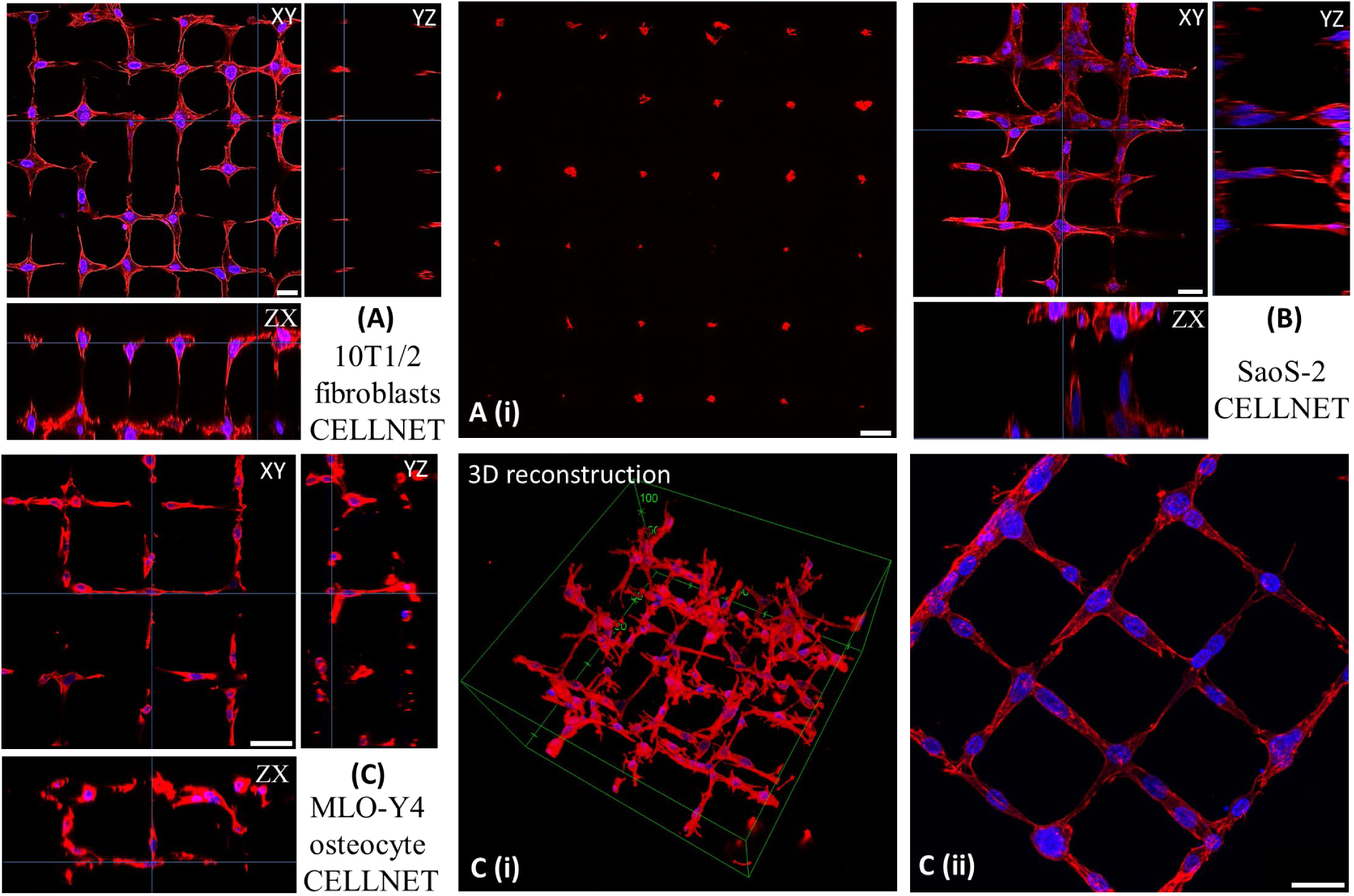
(A-C) Morphology of CELLNETs using three model cells. Cross-sectional image in A(i) shows actin-labelled cellular processes extending between two z-planes ∼50μm apart. 3D image C(i) and high-resolution image (Cii) of MLO-Y4 CELLNET Scale bar: 25μm

**Fig. 6.**
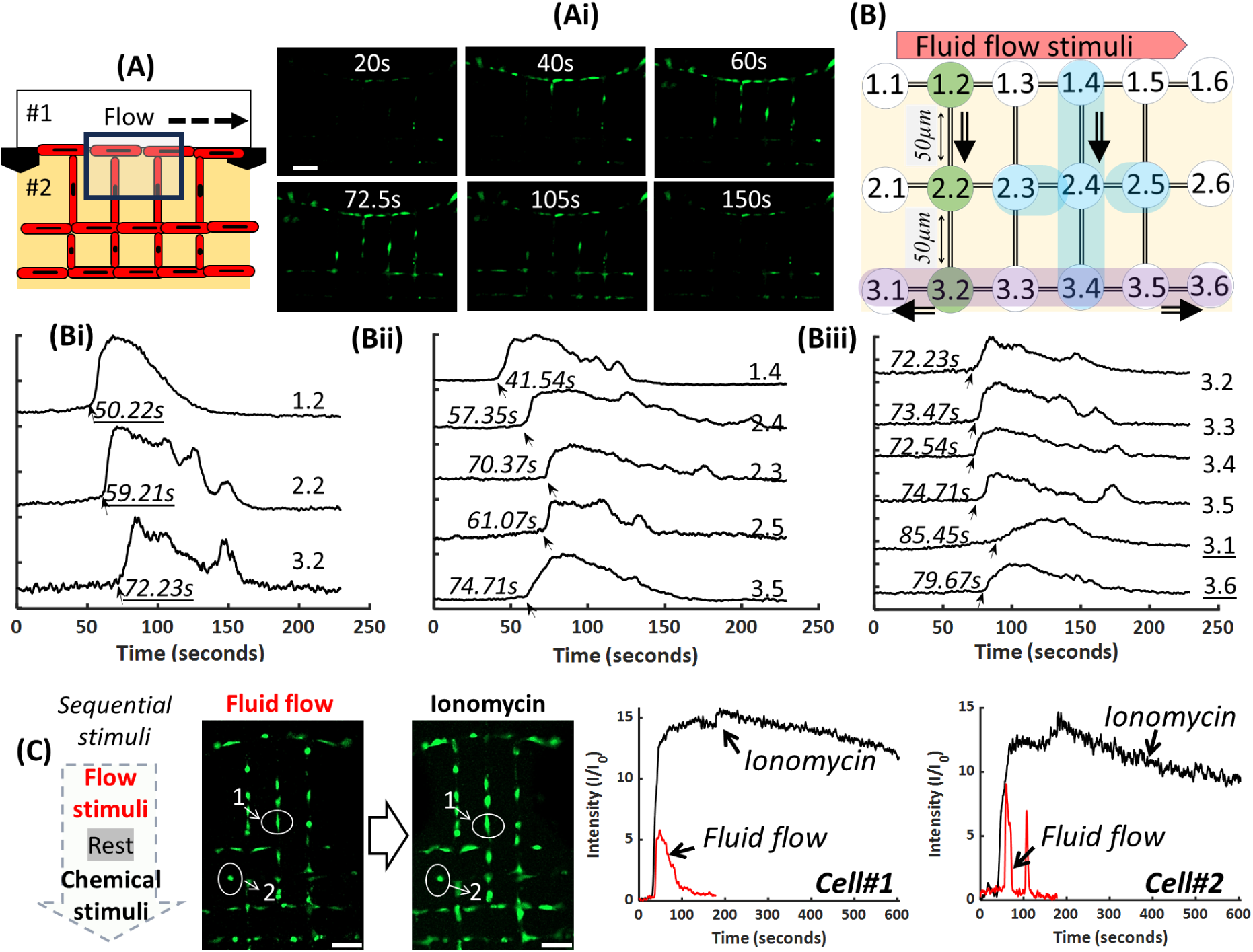
Real-time signaling within osteocyte (MLO-Y4) CELLNETs. **(A)** Schematic showing application of a biophysical fluid flow stimuli in Chamber #1. **(Ai)** Time-lapse fluorescence images showing calcium signaling at specific time-points. **(Aii)** Schematic of model CELLNET. Here, 1.2 represents a single cell in row 1 and column 2, located at the interface of Ch#1-2. Upon stimulation, 1.2 initiates a Ca signal that propagates to adjacent connected cells embedded deeper within collagen (2.2 and 3.2) **(Bi-iii)** Normalized single cell signals from three representative local networks (time-stamps are shown in seconds). **(C)** Signaling response of individual osteocytes (Cell #1 and #2) and the entire network can be accurately analyzed in the presence of sequential biophysical and biochemical stimuli. Scale bar: (Ai), C: 50µm

**Fig. 7.**
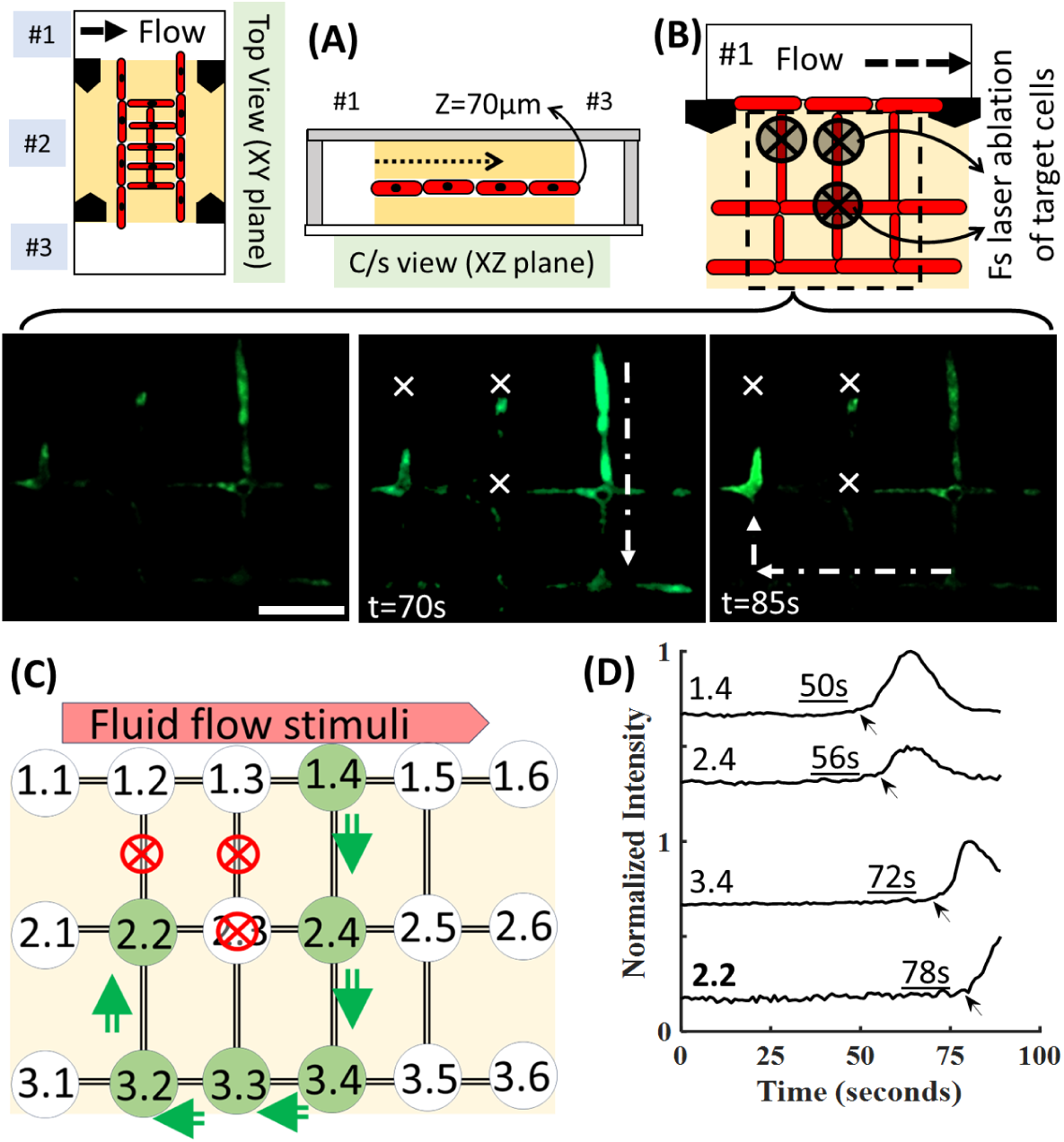
In-plane (∼70µm inside collagen) real-time signaling within user-disrupted osteocyte CELLNETs. **(A,B)** (i) Schematic showing application of a biophysical fluid flow stimuli in Chamber #1 after femtosecond laser ablation was used to lethally injury three cells (shown by cross mark). **(C)** Simplified representation of how signal propagation is diverted due to targeted disruptions. **(D)** Intensity plots showing delayed propagation of signal to cell 2.2 as signal travels from cell 1.4 to 2.4 to 3.3 to 2.2. Scale bar: B: 50µm

### Testing functionality of CELLNETs using stimuli evoked real-time Ca signaling studies

We choose to generate MLO-Y4 CELLNETs to mimic an organized 3D osteocyte interconnected networks found within bone tissue and study the real-time signaling responses of individual cells and the entire network when subjected to biophysical and biochemical stimuli. Before monitoring of real-time signaling within CELLNETs, the viability and morphology of osteocytes were analyzed. Results show an organized square grid network of cells with high viability of 91.39 ± 1.08%.**(SI-Fig.6A)** Osteocytes form 3D in-plane and out-of-plane cell-to-cell connections as evidence by the 3D reconstruction and orthogonal views. As explained earlier, number of cell-to-cell connections were characterized. Specifically, 22.45±2.04% cells connect with 5 neighboring cells (4 in-plane and 1 out-of-plane connections), 20.07±0.59% connect with 4 cells, 25.17±2.36% connect with 3 cells, 11.56±1.56% with 2 cells, 5.44±3.28% with one cell, while 15.31±1.77 were not connected to any neighboring cells. Thus, for this 2-layer design, we found that ∼85% of cells were connected to at least one neighboring cell within the network while maintaining single cell occupancy within ablated channels. Since a lumen size of ∼12µm resulted in many occurrences of multiple cells in the same location within the ablated networks,**(SI-Fig.6B)**, we chose a lumen size of ∼8µm and a square grid network design to test signaling within osteocyte CELLNETs. First, MLO-Y4 CELLNETs were generated in the central chambers (#2) of microfluidic devices and treated with Fluo-4 Ca indicator via media chambers (#1, #3). **(Fig.6A)** Timelapse fluorescence images show an increase in calcium signals in cells proximal to the stimuli, and subsequent transmission of the signals to cells embedded deeper within collagen. **(Fig.6Ai)** To track signaling response of individual osteocytes within CELLNETs, a simplified nomenclature ‘x.y’ was used, where x point to row number and y point to column number. **(Fig.6B)** For instance, row 1 represents osteocytes at the interface of central and side chamber that could directly experience the stimuli, while rows 2 and 3 represent osteocytes embedded at increasing distance within collagen (away from the stimuli). Control over laser scanning path allows the generation of a network with deterministic cell-to-cell connectivity. For instance, a black line represents a direct microchannel connection between cell 1.1 and embedded cell 1.2 while an absence of a black line between cell 1.1 and cell 2.1 indicates that these cells are not connected. For individual cells within the network, calcium signaling, expressed as fold change in fluorescence over baseline, was plotted and time-lags between locally connected osteocyte circuits were analyzed. **(Fig.6Bi-iii)** For instance, signals travel from cell 1.2 to 2.2 to 3.2 (a distance of ∼100µm) with peaks recorded at 50.22s, 59.21s and 72.23s respectively (total time of 22.01s for a ∼100µm resulting in speed of ∼ 4.54µm/s). **(Fig.6Bi)** Each cell within a 3D connected network function as a node, which when connected to other cells in the network, relay stimuli evoked Ca signals. Consider another locally connected network (marked in blue), here a clear time delay is seen as signal travels from cell 1.4 (41.54s) to 2.4 (57.35s) to three connected cells 2.3(70.37s), 3.4 (72.54s) and 2.5 (61.07s). **(Fig.6Bii)** Since all cells are not always present at the nodes with equal spacing between them, the location of each cell was analyzed using FIJI image processor to get accurate measurements about propagation speeds. **(SI-Fig.8)** Lastly, consider all cells in third row; here cells 3.1 and 3.6 are not connected to cells from previous rows, while cells 3.2, 3.3, 3.4 and 3.5 are connected to cells in row 2. As expected, the signal travels to the connected cells first before propagating to 3.1 and 3.6 via 3.2 and 3.5 cell-cell connections respectively. **(Fig.6Biii)**.

**Fig. 8.**
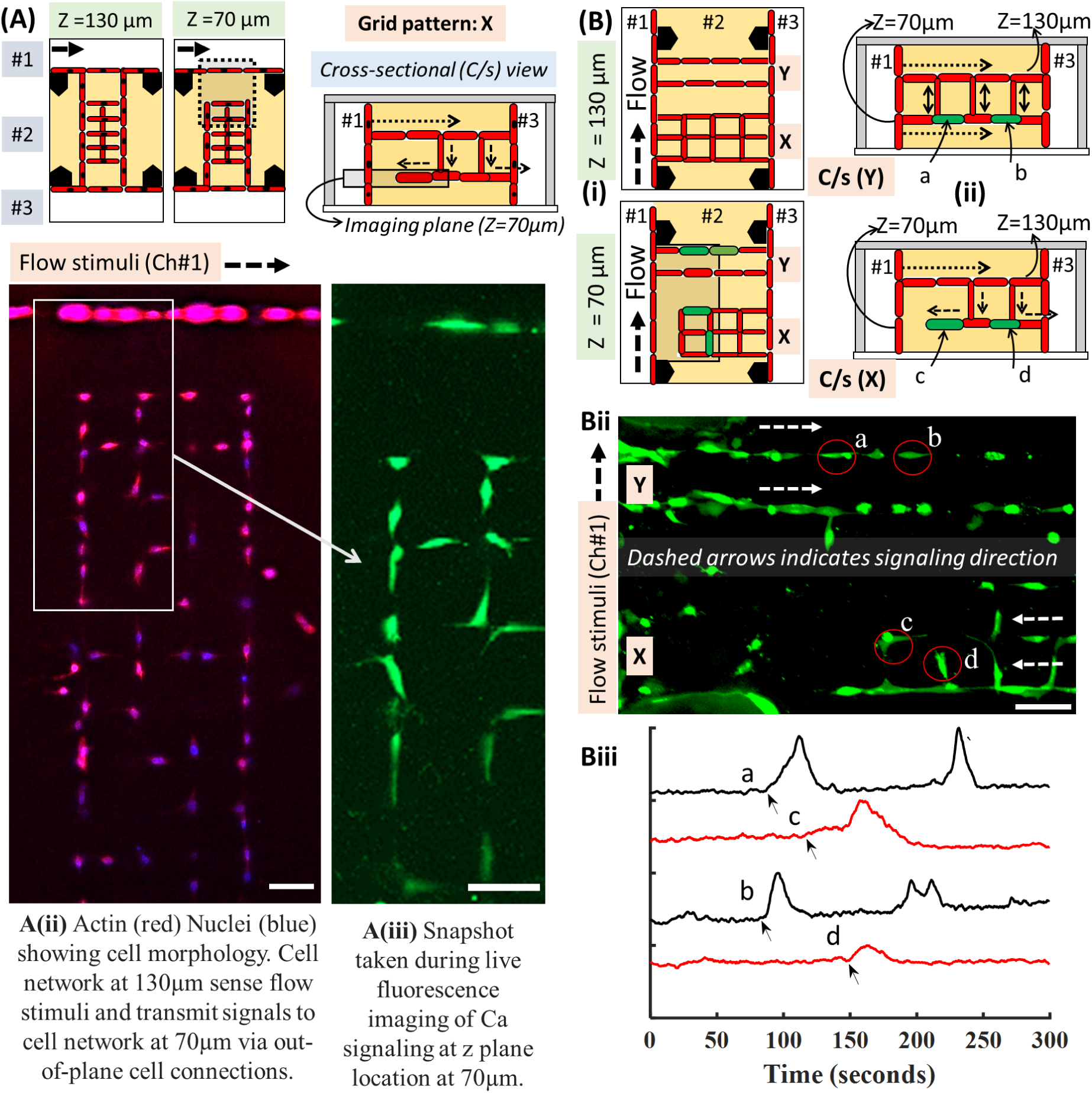
Out-of-plane (3D) real-time signaling within user-disrupted osteocyte CELLNETs. (A) Schematic of 2-layered network with cells organized using a grid patterns (X). Top views (i) and cross-sectional view (ii) showing out-of-plane signal propagation. **(B)** (i) Schematic of 2-layered network with two different cell patterns (X=grid, Y=parallel line) located near each other in the same sample. Top and cross-sectional views of both patterns showing out-of-plane signal propagation **(ii)** Snapshot taken at a z-plane of ∼70µm during calcium signaling experiment. Cells a, b in pattern Y and c, d in pattern X have been marked by a red circle, while dashed white arrows indicate signal propagation directions. **(iii)** Signal intensity peaks of cells ‘c’ and ‘d’ in pattern X are delayed as the signal travels from the cell-network @z=130µm to cell-network @z=70µm as compared to cells ‘a’ and ‘b’ in pattern Y where flow stimuli has a direct path via cell network @z=70µm. Note: In pattern Y, both top and bottom layers are directly exposed to flow stimuli while in pattern X, only the top layer is directly exposed to stimuli.

We then provide a sequential flow stimulus, followed by rest, followed by a chemical stimulant (ionomycin, 5μl, 45μmol.L^-1^, commonly used ionophore known to enhance Ca signals). **(Fig.6C)**. The sequence of fluid flow, rest, and ionomycin stimulus was performed on the same sample, and resulting changes in calcium flux were captured. Here we pick two osteocytes (marked as #1, #2) within the CELLNET and monitored their response over time when the network is subjected to a defined set of sequential stimuli. Results show signaling heterogeneity in magnitude and signal profiles between the two cells and between the stimuli type for the same cell. **SI-Fig.9** shows calcium intensity plots for 15 individual osteocytes within the network. We note that, for all cells within the CELLNET, ionomycin generates a stronger Ca signaling response often lasting for more than 10 minutes as compared to the response evoked by the flow stimulation. Like earlier results, cells closest to the stimuli are triggered first, followed by signal propagation inside the entire network. Time delays within the network is directly related to the user-templated microchannel design and associated cell-to-cell connectivity.

### Study of Ca signaling and signal propagation within custom designed disrupted CELLNETs

Here, we show the CELLNETs can be disrupted in custom configuration by ablating individual cells within the network by targeted femtosecond laser irradiation. First, ablation threshold to lethally injure target osteocytes was identified using via standard live/dead assay. Osteocyte monolayer, labeled with green cell tracker, were identified, and irradiated with femtosecond laser at varying power (10-200mW) and exposure times (0.5-1s). Ablation threshold to cause permanent injury was identified to be 0.22J/cm. To compensate for a decrease in laser dosage while irradiating cells deep in collagen (∼150µm), ablation was performed with 3x of the threshold used under monolayer conditions. Thus, a lethal dosage of 0.64J/cm^2^ (Objective: 40X, NA:0.8, Power: 150mW, exposure time:0.5s) was used for all disruption experiments in this work. Here we tested two network designs.

First design involves single-layer MLO-Y4 CELLNETs in square grid orientation located ∼70µm inside collagen in the central chamber (#2) of the microfluidic device. **(Fig.7A)** On Day 5, focused femtosecond laser was used to lethally injure individual osteocytes. **(Fig.7B)** Then, Fluo-4 Ca indicator treatment was applied and timelapse fluorescence imaging was used in the presence of fluid flow stimuli to monitor real-time Ca signaling within the disrupted CELLNET. Real-time fluorescence images show an absence of Ca signal and propagation for injured cells within the network. Based on the simplified nomenclature described earlier **(Fig.7C, SI-Fig.10)**, cells 1.2, 2.2 and 2.3 were lethally injured forcing the signals to propagate from cell 1.4 (proximal to stimuli) to cell 2.4 (row 2, column 4) to other cells within the network. Plots shows the time-delays of signaling peaks as the signal travels from cell 1.4 (50s) to cell 2.4 (56s) to cell 3.3 (72s) to cell 2.2 (78s). **(Fig.7D)** In an undisrupted CELLNET, signal would have chosen the shortest path to propagate to cell 2.2 (i.e, from cell 1.2 to cell 2.2), however due to user-defined disruptions, the signal is forced to circumnavigate using the only possible connected path available, as indicated by shifts in the signal intensity peaks and associated time-delays.

Design 2 involves a 2-layer CELLNET in a square grid orientation indicated by ‘X’ in **Fig.8**, where the top and bottom layers are located at a depth of 130µm and 70µm inside collagen respectively. For pattern ‘X’, only cell network located at z=130µm has direct physical connection to chamber #1 while the cell network located at z=70µm can only indirectly sense the stimuli through out-of-plane cell-to-cell connections. Fluorescence images show osteocyte morphology **(Fig.8Aii)** and a snapshot of Ca signaling **(Fig.8Aiii)** from the imaging plane located at z=70µm. These results clearly show that there is no direct contact between the flow stimuli applied in Ch#1 and cell network located at z=70µm.

To compare signal propagation speeds between directly and indirectly connected cell networks, we prepared two network 3D patterns ‘X’ and ‘Y’. **(Fig.8Bi)** While both patterns are located at z=130µm and z=70µm inside collagen, for parallel-line pattern Y, both top and bottom cell networks can directly sense flow-stimuli evoked signaling response while for pattern ‘X’, cells located at z=70µm can indirectly receive signals from stimuli-evoked cells located at z=130µm. Fluorescence image showing a snapshot of Ca signaling within the cell network located at z=70µm. **(Fig.8Bii, SI-Fig.11)** (Arrows indicate the signaling direction for both patterns). Cells ‘a’ and ‘b’ from pattern Y and cells ‘c’ and ‘d’ from pattern X are chosen as representative cells, and their signaling responses are compared when subjected to a flow stimulus. **(Fig.8Biii)** Peak delays observed in ‘c’ and ‘d’ as compared to ‘a’ and ‘b’ indicate that pattern X enforces Ca signal propagation from the top-layer (z=130µm that directly senses the stimuli applied in Ch#1) to cells in the bottom layer (z=70µm). These results show the capability of studying real-time signaling within user defined disrupted cellular networks.

## Discussion

All tissues are composed of single cells, yet how tissues sense, respond, and adapt to stimuli cannot be predicted by studying individual cells as higher-level emergent property of the tissue likely rely on complex dynamic interactions between cells. To do this, the field needs new ways to generate tissue-specific 3D single cell networks, apply defined stimuli, and analyze networks’ adaptations. At present, such a technology does not exist. As a result, the link between the individual cells and cell-networks’ (tissue) function remains unclear and difficult to predict. Here, we report a novel technology, coined as CELLNET, to generate normal and disrupted 3D single-cell networks within type I collagen matrix in user-defined configurations and study the real-time signaling of single cells and signal propagation across the entire network. We show that this template-based strategy works with many cell types, is highly reproducible and provides user control over cell-cell connectivity and cell-network layout. Use of DLP allows rapid and inexpensive iteration of master molds enabling rapid production of custom internal microchannel designs within standard microfluidic devices. We show that real-time Ca signaling of individual cells and signal propagation within CELLNETs can be monitored when subjected to biophysical and biochemical stimuli. Moreover, femtosecond laser irradiation can ablate target cells within CELLNETs at defined locations and time-points to design custom disrupted networks and study real-time changes in their signaling dynamics. This allows entire CELLNETs or individual cells within the network to be manipulated in a noninvasively, contactless, and sterile manner during active culture.

To test the capability of studying real-time signaling within CELLNETs, we choose osteocytes as our model cells, due to our groups’ prior experience with bone tissue engineering.^30^ Like many tissues, in bone, stimuli evoke Ca^2+^ signals within 3D, organized, single-cell osteocyte networks while signal disruptions have been linked to many pathologies. Similar to other cell types, single-cell osteocyte networks have also been generated using micropatterned cell-adhesive self-assembled monolayers (SAMs) or micro-chambers^31–32^, or two-photon laser based modifications^9, 28^, however generating single-cell network in 3D remain difficult.^33–37^ As a result, culture cells within collagen or fibrin matrices remain the ‘gold standard’ to generate 3D cell networks, however the randomly organized cell-cell connections with poor reproducibility makes a systematic study about single cell and its relationship to network signaling difficult^38–49^. In contrast, structurally defined 3D MLO-Y4 (osteocyte) CELLNETs simplify image processing and enable accurate mapping of how single cells respond to various types of stimuli and their contributions to changes in signaling wavefronts within the networks.

Previously, our group and others have used femtosecond laser ablation to modify local properties within a 3D cell-laden hydrogel matrix to guide the alignment of single cells. For instance, our group showed laser based densification of partially crosslinked GelMA was used as guidance cues to align encapsulated cells in 3D^15^. Another study used modified PEGDA hydrogels to generate cell network.^28^ In both these cases, native extracellular matrix like collagen cannot be used due to the requirement of the photo-sensitive hydrogels to enable laser-based biochemical or biophysical modifications. Both laser processing conditions and hydrogel properties must be optimized to maximize viability of encapsulated cells; this enforces strict constraints on hydrogel type and processing conditions and limits its utility in the field. Laser scanning also generates reactive oxygen species (ROS) which decrease cell viability by interfering with its metabolism. Since ROS generation is strongly dependent on applied laser dosage and the material photosensitivity, new semi-synthetic hydrogels with better photosensitivity are needed to satisfy the contrasting requirements of rapid hydrogel modifications (either ablation^7, 9^ or degradation^6, 8^) while maintaining high cell viability. With CELLNET, a user-defined templated in collagen is generated before cells are seeded/pipetted in target microfluidic chambers. This decoupling provides flexibility to generate 3D cell networks in any bioink including native and unmodified ECM like collagen which in turn results in close to 100% cell viability as laser scanning is not performed in the presence of cells. In this work, we use type I collagen (4mg/ml) as our model ECM as this natural and thermally crosslinkable biomaterial has been widely used in the field. We have also tested other ECM analogs such as bovine collagen and semi-synthetic PEGDA-GelMA hydrogels, as seen from **SI-Fig.12**. We envision CELLNET to be an ECM- and cell-agnostic technology that can be broadly used to design tissue-specific CELLNETs for a range of applications.

CELLNET could also emerge as an ideal tool to study signaling heterogeneity, a potentially important yet unstudied phenomenon. Previously, we and others have shown that self-assembled and randomly connected osteocyte networks exhibit spatial variation and temporal heterogeneity of Ca^2+^ waveforms when subjected to fluid flow stimuli.^50^ Due to difficulty in identifying individual cells in self-assembled and randomly organized networks, the relationship between cell connectivity, stimulus conditions, and signal heterogeneity cannot be studied. In this work we use calcium signal as a proxy to monitor real-time signaling of individual osteocytes within spatially organized 3D single cell networks. CELLNETs within microfluidic devices provides systematic application of biophysical, biochemical or injury stimuli that could be used to assess single cell signaling heterogeneity within other cell types.

Overall, the modular nature of CELLNET allows ‘swapping in’ relevant cell types, ECM composition, network designs, optical tracers for other signals (NO, ROS, ATP) and/or fluorescent reporters or inhibitor drugs and acquire new knowledge using standard imaging and culture methods. In the future, transfected variants of cell lines with genetically encoded calcium indicators can facilitate fluorescence imaging without exogenous labels and allow us to study the dynamic evolution of CELLNETs over time. The simple and easy to use cell seeding strategy to generate CELLNETs will enable broad utility and adoption in the field to test new hypothesis across cell, tissue, or stimulus types and develop personalized tissue-specific models for drug testing and diagnostic applications. We envision that CELLNET will transform the study of tissue-scale system biology by linking individual cells and tissue networks’ function thereby elucidating higher-level emergent property.

## Materials and Methods

### Fabrication of PDMS microfluidic devices using DLP-printed PEGDA Master Molds

Prepolymer solution was prepared using poly(ethylene glycol) diacrylate (PEGDA, MW= 250Da; Sigma-Aldrich) as the base material, with Phenylbis(2,4,6-trimethylbenzoyl)phosphine oxide (commonly known as Irgacure 819; Sigma-Aldrich) and 2-Isopropylthioxanthone (ITX; Tokyo Chemical Industry) as the photoinitiator and photosensitizer respectively, while 2,2,6,6-tetramethyl-1-piperidinyloxy (TEMPO; Sigma-Aldrich) served as the free radical quencher. A stock solution was prepared by mixing 100 mg of Irgacure 819, 200 mg of ITX, and 4 mg of TEMPO with 40 ml of PEGDA in a centrifuge tube, which was then wrapped in aluminum foil and vortexed to ensure thorough mixing of the chemicals in the solution. This stock solution was then stored at room temperature until use.

Prior to printing, the glass slides were subjected to cleaning in piranha solution (H_2_SO_4_ and H_2_O_2_; 7:3) with constant stirring at 125 rev/min for 30 mins, followed by rinsing with ethanol and water and subsequent drying at 65 °C in a vacuum oven for an hour. The surfaces of the glass slides were then modified with methacrylate groups by immersing them in a solution containing 3-(trimethoxysilyl)propyl methacrylate (TMSPMA; Sigma-Aldrich) and Toluene (Sigma-Aldrich) (9:1) at 50°C, while maintaining a constant stirring at 125 rev/min. The acrylated glass slides were dried at 65°C in a vacuum oven overnight, and the modified cover slips were affixed to an aluminum print head using a double-sided tape for printing.

Custom Digital Light Projection (DLP) platform was designed and built in Soman group. **(SI-Fig.1A)** Briefly, the optical setup consists of a 405 nm CW laser light source (iBEAM SMART 405, Toptica Photonics), DLP development kit (DLP 1080p 9500 UV, Texas Instruments, USA), and a Z-stage (25 mm Compact Motorized Translation Stage, ThorLabs). A rotating diffuser was added to the setup to eliminate laser speckles generated by the Gaussian beam profile, and to convert Gaussian intensity profile into a uniform hat-shaped distribution beam profile before expanding, collimating, and projecting the beam onto the DMD which consists of an array of 1920 x 1080 micromirrors with a single pixel resolution of around 10 µm. A 3D CAD model of positive master mold of a three-chambered microfluidic device was designed in SolidWorks **(SI-Fig.1B)**. Custom MATLAB code was used to slice the model and generate a stack of binary portable network graphics (PNG) image files. These files were uploaded onto DMD and converted to virtual masks to spatially modulate the laser beam onto a vat of liquid prepolymer solution through an oxygen permeable PDMS window. LabVIEW code was used to control DMD and synchronize it with the upward movement of the print head to photo-polymerize the master mold onto a surface-modified glass coverslip.

PEGDA master molds were used to make the final PDMS microfluidic devices. First, polydimethylsiloxane (PDMS, Sylgard 184: Dow Corning Corporation) base and curing agent were thoroughly mixed in a 10:1 mass ratio, degassed under vacuum, and poured onto the master molds. The molds were initially placed in a 52 mm petri dish, then PDMS was poured directly over the micro holes (reverse feature for micro-pillars) and degassed to ensure uniform filling of PDMS in all negative spaces of the mold. PDMS-mold setup was then cured for 10-12 hours at a low temperature (35-40°C) and for an additional 2 hours at a higher temperature (65°C). After cooling, the PDMS layer was gently peeled from the molds and trimmed to size. Inlet and outlet holes were punched out of the PDMS to form 3 inlets and 3 outlet ports. Finally, the PDMS was irreversibly bonded to a glass coverslip (0.15mm thick) using oxygen plasma treatment (60-seconds exposure).

### Generation of user-defined microchannels within crosslinked collagen

Type I collagen solutions at varying concentrations (Rat tail tendon, ibidi, Germany) were prepared using established protocols. Briefly, stock solution of collagen (10 mg/ml) was diluted to 4 mg/ml with 1X medium (DMEM, ThermoFisher) and 10X medium (HBSS, ThermoFisher) such that the final salt concentration is 1X and it is neutralized to pH 7.2–7.4 by adding calculated amount of 1M NaOH. All the reagents, media and collagen were placed in ice-bath throughout the neutralization process. Final pH was measured using pH strips. Before pipetting the collagen solution into the PDMS microfluidic devices, the channel surfaces were modified with (3-aminopropyl)triethoxysilane (APTES) and glutaraldehyde (GA) to immobilize the collagen and prevent detachment from channel surfaces. First, devices were sequentially immersed in a solution of (i) 10% (APTES, Sigma-Aldrich) in ethanol for 1 hour, followed by three ethanol rinses, and (ii) 2.5% glutaraldehyde (GA, Sigma-Aldrich) solution in deionized water for 1 hour, followed by three rinses using deionized water, and (iii) sterilization under UV light overnight. Collagen solution was pipetted into chamber #2 of the devices, followed by incubation at 37°C for 20 minutes to allow gelation of the matrix. Then, 1X PBS supplemented with 1% Penicillin-Streptomycin (10,000 U/mL, Thermo Fisher Scientific) was pipetted into chambers #1 and #2 to prevent crosslinked collagen from dehydration and bacterial contamination. The devices were stored in incubator for 24 hours to ensure robust crosslinking of collagen and consistent channel sizes during laser ablation experiments. All femtosecond laser ablation experiments were performed after 24hours at user-defined time-points.

Custom-built fs-laser ablation setup was designed and built by combining a Ti:Sapphire fs laser (Coherent, Chameleon, USA) with a Zeiss Microscope (Observer Z1, Germany). **(SI-Fig.2A)** In this setup, an 800 nm femtosecond laser beam with a repetition rate of 80 MHz was focused inside crosslinked collagen within the microfluidic devices using a water immersion objective (40×, NA=0.8, Leica). User-defined patterns, designed using visual basic code, were used to scan the laser focus within the collagen to ablate 3D microchannels. To control the microscope stage, we developed a Visual Basic script that was integrated with Axio Vision software (Zeiss, Germany) to precisely manipulate the XYZ stage to trace complex trajectories. For example, to create a helical channel, we determined the locus of points (x, y, z) from online resources and used it to write a visual basic code to automate microscope stage movement in 3D. A custom written algorithm was used to synchronize the stage and laser shutter to write 3D patterns at controlled speeds and laser powers. The dosage was changed by modulating the average power of the laser using a polarization-based power-tuning system or by modulating the scanning speed of the stage. Ablated patterns in both lateral and vertical directions were visualized in real time under bright-field microscopy. **(SI-Fig.3A-B, SI-Videos 1 and 2)**

Laser ablation of other types of hydrogel matrix was also performed. Bovine collagen (Advanced Biomatrix, 7mg/ml) was used similarly to the rat tail collagen described earlier. Additionally, semi-synthetic gelatin methacrylate (GelMA, Advanced Biomatrix), synthetic polyethylene glycol diacrylate (PEGDA, 6KDa, Advanced Biomatrix), and their combinations were tested. Here hydrogel solutions were pipetted in Ch#2 and crosslinked for 15 seconds using LED light Cure Box (B9A-LCB-010: B9Creations) that emits light at 390–410. Microchannels were then created using femtosecond laser ablation. Here, Lithium Phenyl (2,4,6-Trimethylbenzoyl) Phosphinate (LAP; Signa-Aldrich) was used as photoinitiator.

### Characterization of 3D cell networks within target chambers of microfluidic devices

<u>Murine osteocyte-like cell line MLO-Y4</u> (Kerafast, Inc. Boston, MA) was cultured according to the suppliers recommended protocol. Briefly, T75 vented cell culture flasks were coated with 4 μg cm^−2^ rat tail type I collagen (Sigma-Aldrich, Inc. St. Louis, MO) for 30 minutes at 37°C before cell seeding. MLO-Y4 cells were cultured in these coated flasks using minimum essential media (α-MEM, GIBCO#12571-063) containing L-glutamine, Ribonucleoside and deoxyribonucleosides; supplemented with 2.5 % heat inactivated Fetal Bovine Serum (FBS, R&D Systems, Minneapolis, MN), 2.5% calf serum (Cytiva Life Sciences, Marlborough, MA) and 1% penicillin/streptomycin (Thermo-Fisher Scientific). <u>The fibroblast-like 10T1/2 cell line</u>, derived from a C3H mouse embryo cell and exhibiting fibroblast morphology, was bought from American Type Culture Collection (ATCC, Manassas, VA) and cultured in T75 vented culture flasks using a cell growth media consisting of a Basal Medium Eagle (BME, GIBCO#21010-046) supplemented with 10% fetal bovine serum (FBS), 1% penicillin/streptomycin and 1% GlutaMAX (GIBCO#35050-061). <u>MC3T3-E1, pre-osteoblast cells</u>, provided by Dr. Horton (SUNY Upstate Medical University), were grown in α-MEM (GIBCO#A1049001) supplemented with 1% GlutaMAX, 1% sodium pyruvate, 1% penicillin/streptomycin and 10% heat inactivated Fetal Bovine Serum. <u>Saos-2</u> <u>(ATCC, Manassas, VA), human osteosarcoma cell line</u>, was cultured in Dulbecco’s modification of Eagle’s medium (DMEM, Gibco#11965-092) as the foundational medium, supplemented with 10% heat-inactivated fetal bovine serum (FBS), 1% GlutaMAX and 1% penicillin-streptomycin. All the cells were maintained in a controlled environment at 37°C with 5% CO_2_ and 90% humidity.

For microfluidic devices, all the cell types were harvested by trypsin treatment (0.05 × 10^−3^ M in 1X PBS, 5 min), centrifugation, and media washing. Cell seeding was performed by pipetting of harvested cells (1 × 10^6^ cells/ml) into the inlet ports of one of the side chambers (#1 or #3). Capillary action and differential pressure facilitate the perfusion of cell solution within the microchambers as well as the 3D ablated microchannel networks. Cells were allowed to settle in the network for 30 mins before flushing non-adherent cells from the device using corresponding growth media, and devices were cultured in 35mm cell culture petri dish under static culture conditions (37°C, 5% CO_2_ and 90% humidity). Daily monitoring of cell growth using brightfield microscopy was conducted, with media channels being flushed each time cells were observed, and periodic brightfield images were captured. **(SI-Fig.4A,B)** If necessary, media channels were flushed with trypsin to prevent unwanted cell accumulation in the side chambers. For heterotypic cell seeding, MC3T3 cells were tagged with red CM-Dil dye (Invitrogen#C7001) and MLO-Y4 cells were tagged with either red CM-Dil dye or green CMFDA dye (Invitrogen#C2925) as required. **(SI-Fig.7)** Briefly, cells cultured in T75 vented flasks were harvested and incubated in the corresponding culture media containing 2μg/ml of cell tracker dye for 30 minutes in 1.5 ml centrifuge tubes. Cells were then diluted by adding 5 ml PBS and collected by centrifuging the cell solution. Tagged cells were then seeded in channel #1 and channel #3.

For assessing cell viability, an Invitrogen Live/Dead assay kit, comprising Calcein AM and Ethidium homodimer, was utilized. Briefly, a solution containing 0.5 μl/ml of Calcein AM and 1 μl/ml of Ethidium homodimer in phenol red-free cell media (FluoroBrite, DMEM, GIBCO#A18967-01) was introduced into chambers #1 and #3. The devices were then incubated at 37°C for an hour, and images at different planes were captured using an inverted microscope (Nikon Eclipse). The captured images were processed using FIJI. Cells in two-layered connected square grid structures with 9 x 9 nodes for 10T1/2 cells and 7 x 8 nodes for MLO-Y4 cells were analyzed. Error bars were calculated as standard deviation of data collected from 3 different samples (n=3) for each cell types. To study cellular alignment, the cells were fluorescently stained for f-actin and nuclei and imaged under LSM 980 confocal microscope (Zeiss, Germany). First, 4% formaldehyde (Sigma-Aldrich) was pipetted in Ch#1 and #3 for 30 minutes at room temperature and washed three times with 1X PBS. Cells within Ch#2 of the device were permeabilized with 0.5% TritonX-100 in 1X PBS for 30 minutes. Subsequently, the cells were stained with rhodamine-phalloidin (Invitrogen) at a dilution of 1/250 in 10% horse serum for 45 minutes at room temperature to visualize f-actin, and DAPI (ThermoFisher) at a dilution of 1/1000 for 5 minutes at room temperature to visualize cell nuclei. Samples were washed subsequently washed and stored in 35 mm petri dish covered with aluminum foil covered to protected from light and prevent bleaching of fluorophores. 3D images of CELLNETs in connected two-layered square grid were acquired using confocal microscope (Zeiss LSM 980, Germany) and processed with FIJI image processing software. Grid of 8 x 8 nodes (area: 400μm^2^ in each plane) in the CELLNET was analyzed. The total number of cells in the CELLNET were determined by manually counting the number of nuclei present in the nodes. In the CELLNET with ablated channel size of 8μm, mostly had one cell was present per node, whereas for the channel sizes of 12-15μm, there were multiple cells per node. To access connectivity of cells, stack of f-actin-stained images were analyzed. The analysis was performed in two steps. First, cell connectivity in a single plane was analyzed by taking a single cell in a node and checking if its dendritic processes were connected to adjacent cells. In a two-layered network, single cells in are particular plane could potentially connect to a maximum of four cells (4 in-plane connections and 1 out-of-plane connection). Second, the z-stacked images were resliced to obtain images in the X-Z plane to assess the connectivity of a particular cell to out-of-plane cell connections (z-planes). The newly acquired stacked images were analyzed frame-by-frame to check the connections of cells in different z-planes. Cell connectivity is presented as the percentage of cells connected to the number of adjacent cells in the CELLNET. Error bars represent the standard deviation of data acquired from 5 different samples (n=5, 10 grids and 640 nodes).

### Real-time calcium signaling within ‘deterministic’ 3D single-cell network

Calcium Signaling Fluo-4 Calcium Imaging Kit (Invitrogen#F10489) dye was used to conduct calcium signaling experiments. Prior to mechanical loading or chemical stimulation, Fluo-4 calcium reporting dye was pipetted into side chambers of the device to enable diffusion mediated loading of dye into MLO-Y4 CELLNETs (3D cell networks in collagen in Ch#2). Briefly, the Fluo-4 AM loading solution was prepared by mixing 10 parts of 100X PowerLoad and 1 part of 1000X Fluo-4, AM, with 1000 parts of cell medium. Next, the cell medium was removed from all inlets and oulets of the device, and side chambers (#1 and #3) were gently rinsed with 1X PBS (37°C). 100µL of Fluo-4 AM loading solution was added to both inlets and incubated at 37°C for 30 minutes, followed by 15 minutes at room temperature. Fluorescence data were collected at 1-3 frames per seconds under mechanical and/or chemical stimulation using confocal microscope (Zeiss LSM980) equipped with enclosed chamber maintaining cell culture conditions. <u>Mechanical</u> <u>stimulation.</u> Fluo-4 AM loading solution was removed from the devices. Next, microfluidic chips were mounted on the microscope stage for an additional 10 minutes allowing cells to settle down; this avoids recording of unwanted cell signals due to perturbations during the media removal steps. The data acquisition was started 30 seconds prior to applying mechanical stimulation to the cells. To apply mechanical stimulation to the cells, 150μl of fresh media was pipetted in the inlet of Ch#1. This causes the media to flow from the inlet to the outlet of Ch#1, generating shear stress at the interface of Ch#1 and Ch#2 which is sensed by the cell network located at different z-planes in collagen matrix. Fluorescence signal (changes in calcium within each cell of the network) were recorded using time-lapse fluorescence microscopy, and relative changes in fluorescence intensity of each cell in the network was calculated using established methods. Briefly, calcium signal intensity was quantified with FIJI. Each cell in the 3D network was selected for intensity measurements. Intensity of first 30 seconds, prior to any stimulation, were averaged to generate an initial baseline intensity I_0_. Then the intensity of each frame was divided by the baseline intensity to get fold increase of intensity I/I_0_. All the experiment were repeated at-least three times and the time lag between the signal transfer from one cell to another according to their location in the circuit was analyzed using Microsoft Excel and MATLAB. <u>Biochemical stimuli.</u> For biochemical induced cell responses, 5μl of ionomycin (45μmol.L^-1^) (Invitrogen#I24222) was introduced into the reservoirs containing 180μl of media, post baseline image acquisition, and confocal microscopy was used to record series of images for 10 minutes. Control experiments with 5μl of media was pipetted into Ch#1 of the devices to confirm that mechanical forces during pipetting does not generate any mechanical stimuli and associated calcium signaling responses; results showed no change in the signal intensity of loaded Fluo-4 AM. To model the effects of network discontinuity, femtosecond laser irradiation a dosage of 0.64J/cm^2^ was used; this is calculated based on a 40x objective, 0.8 NA, an aperture size of 9.3mm with power of 150mW and an exposure time of 0.5s. Femtosecond laser was used to lethally injure individual osteocytes within MLO-Y4 CELLNETs and resulting changes to signal propagation and time-delays were analyzed.

## Acknowledgments

We thank J. Horton (SUNY Upstate Medical University) for providing us with MC3T3 pre-osteoblast cells, and assistance with the review of this manuscript. We would like to thank Zachary Geffert for expanding and maintenance of cell lines, and Minh Thanh from Dr. Alison Patteson’s lab in the Physics Department for assistance with the confocal reflectance microscope.

## Author contributions

Each author’s contribution(s) to the paper should be listed (we suggest following the CRediT model with each CRediT role given its own line. No punctuation in the initials.

Examples:

Conceptualization: AP, PS, PK Methodology: AP, PK Investigation: AP, UA Visualization: AP, UA, ABM Supervision: PS, PK Writing—original draft: AP, PS

Writing—review & editing: AP, PS, PK, UA, ABM

## Data and materials availability

All data are available in the main text or the supplementary materials.

## Supplementary Materials

**SI-FIG. 1.**
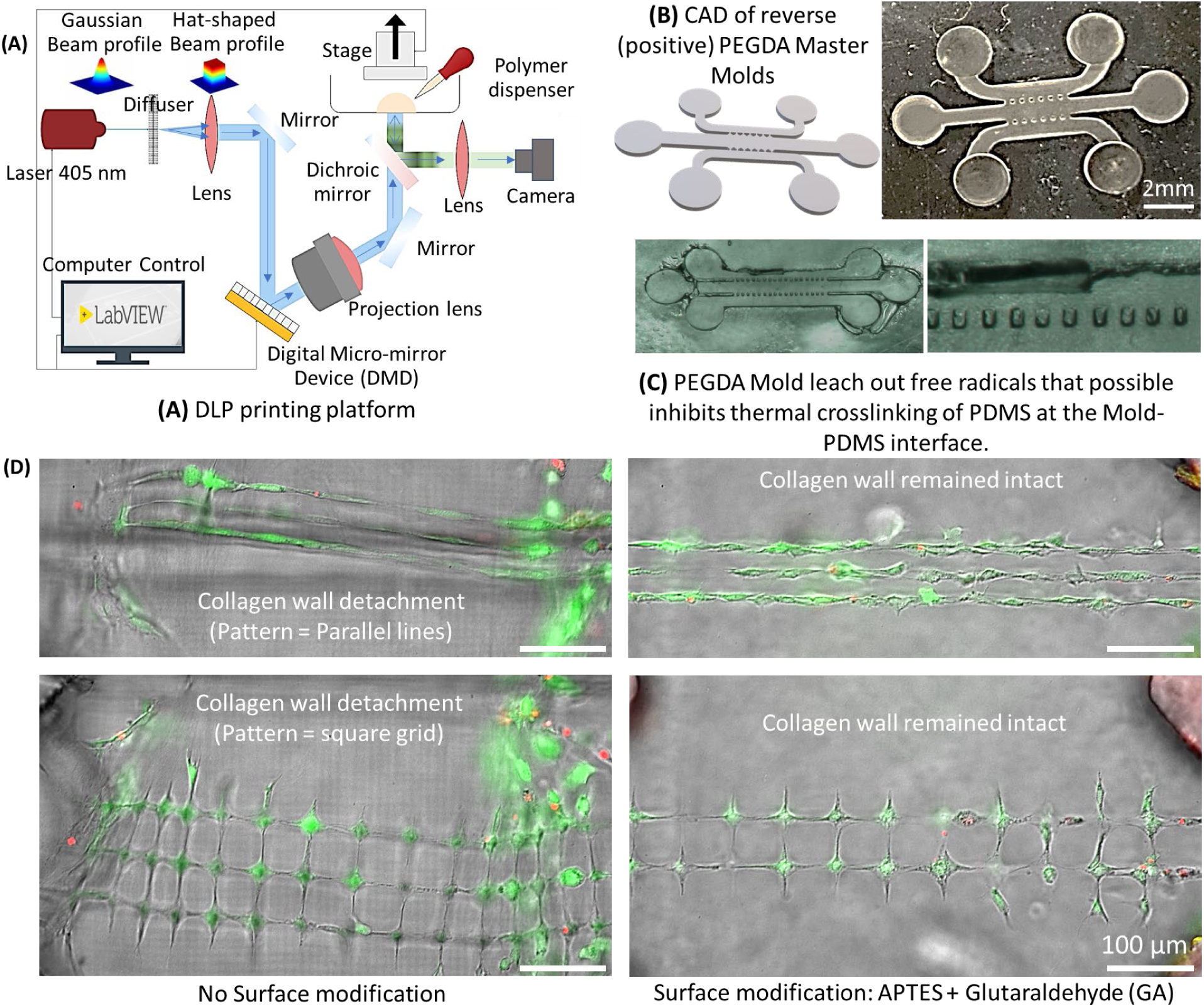
(A) Schematic of Digital Light Projection (DLP) setup to print Master Molds for making multi-chambered PDMS devices. (B) Design files and a representative picture of the Master Molds made using polyethyelene glycol diacrylate (PEGDA 250MW). (C) Pictures of defective molding process in the absence of post-processing steps of ethanol treatment and UV exposure for 24 hours. (D) Composite brightfield-fluorescence image showing parallel line pattern (top row) and square grid pattern (bottom row) with and without surface modification of PDMS-collagen devices. Without surface modifications, crosslinked collagen detaches from the PDMS roof and glass bottom surfaces. Scale bar: 100μm.

**SI-FIG. 2.**
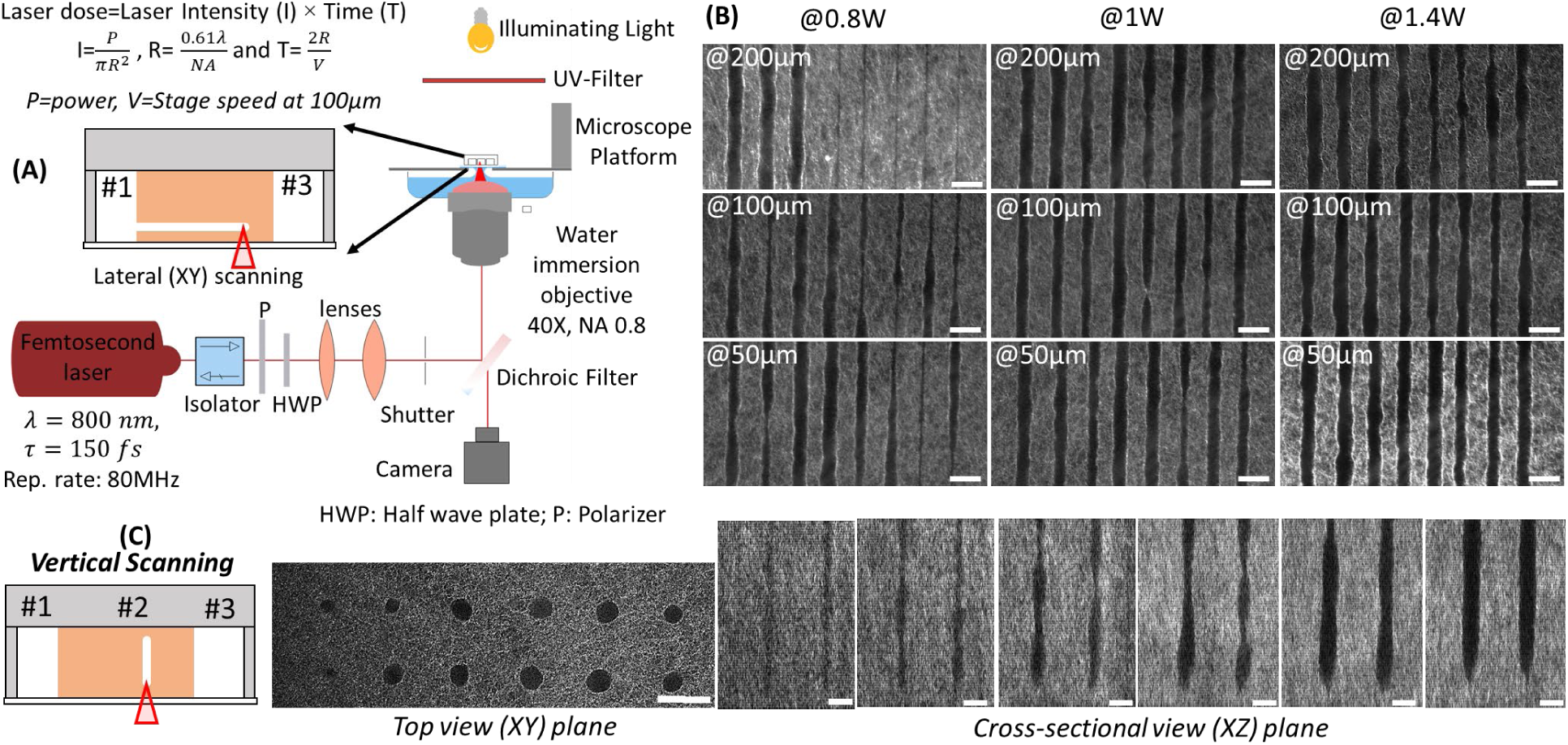
(A) Schematic of two-photon ablation setup. (B) Confocal reflectance microscopy images of ablated channels in lateral (XY) directions at varying powers (0.8W, 1W, 1.4W) and at varying depths (50µm, 100µm, 200µm inside collagen from the bottom glass coverslip). (C) Confocal reflectance microscopy images of ablated channels in vertical (Z) directions under varying conditions. Note that the laser dose is calculated from power of laser right before objective lens and the scanning speed of the stage. (Scale bar: 20µm).

**SI-FIG. 3.**
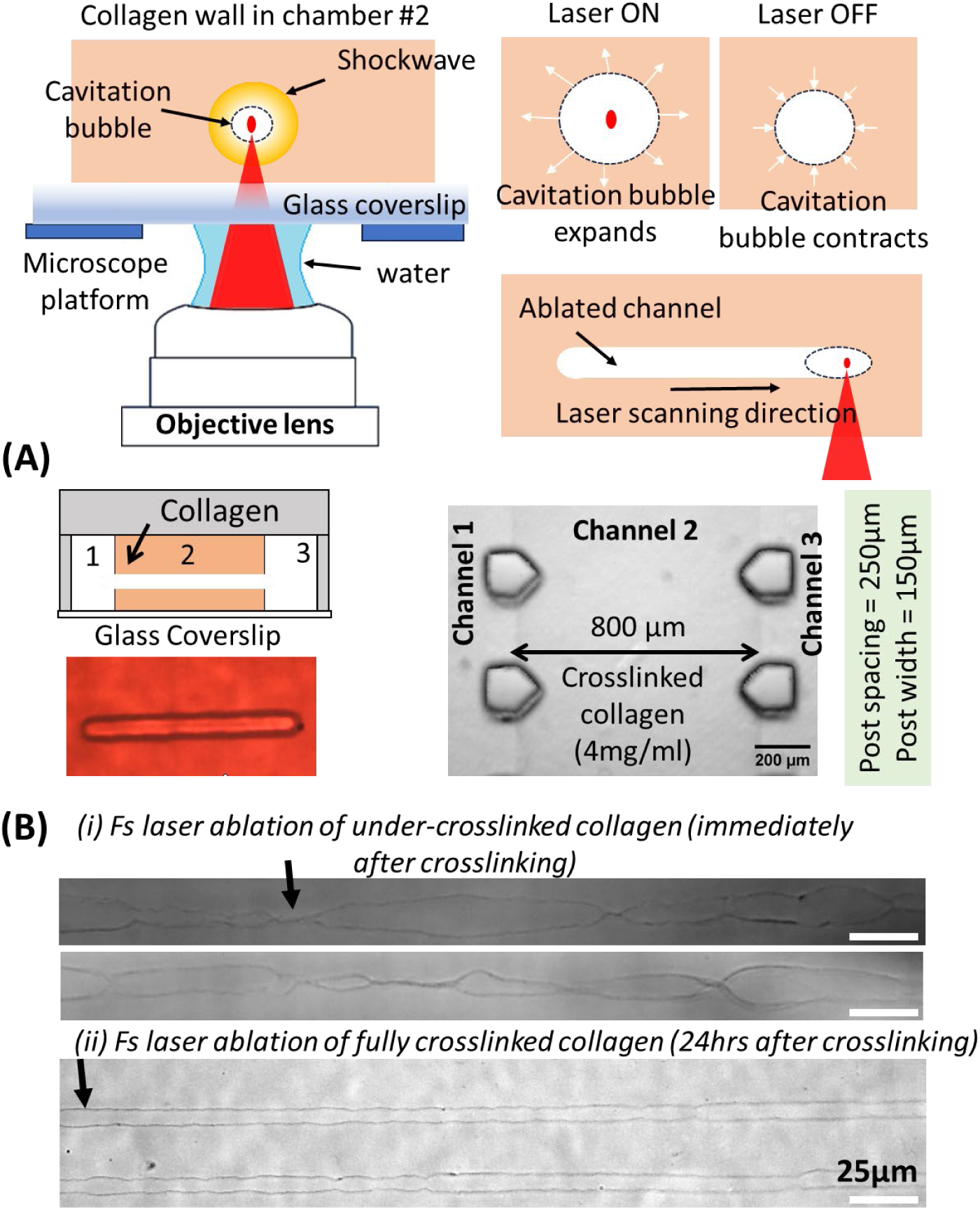
(A) Schematics for laser ablation inside collagen via cavitation. Representative brightfield image of ablated channel in shown (Red color is due to the filter used during imaging). Image of crosslinked collagen in Ch#2 between a pair of micropost arrays is shown. (C) Brightfield images of ablated lines immediately after crosslinking (∼1hr) show multiple location of low fidelity of channel size and shape as compared to ablation performed 24hs after crosslinking under identical laser conditions.

**SI-FIG. 4.**
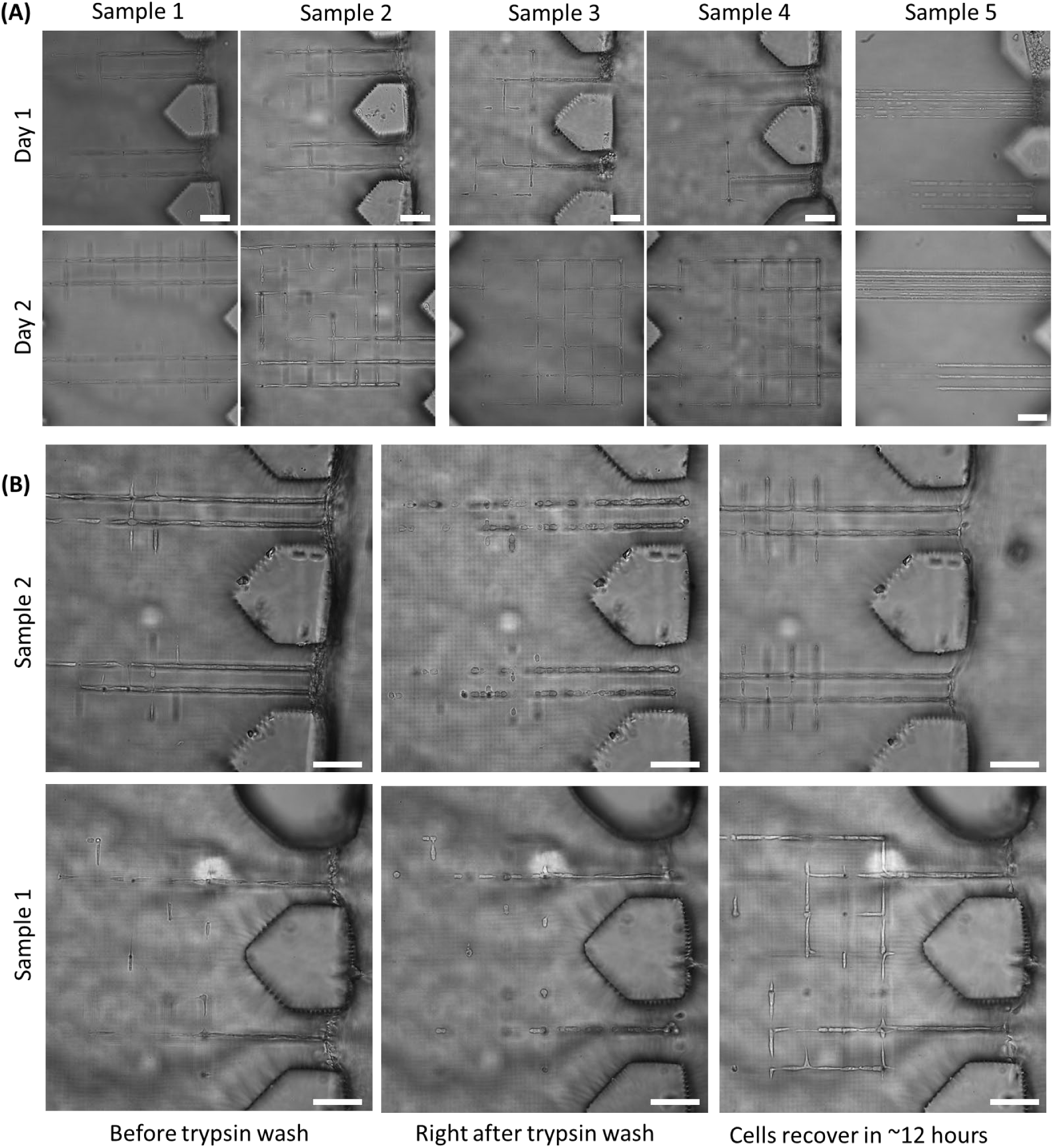
(A) Brightfield images in 5 different samples showing fibroblast migration within ablation microchannel networks generated within crosslinked collagen in Ch#2 of PDMS device. Parallel line and square grid networks are shown. (B) Representative images from 2 different samples showing clearance of cells from the side wall through trypsin treatment (0.25%, 5 mins) and prevent unwanted aggregation of cells in Ch#1 and Ch#3. Scale bar: 100 μm

**SI-FIG. 5.**
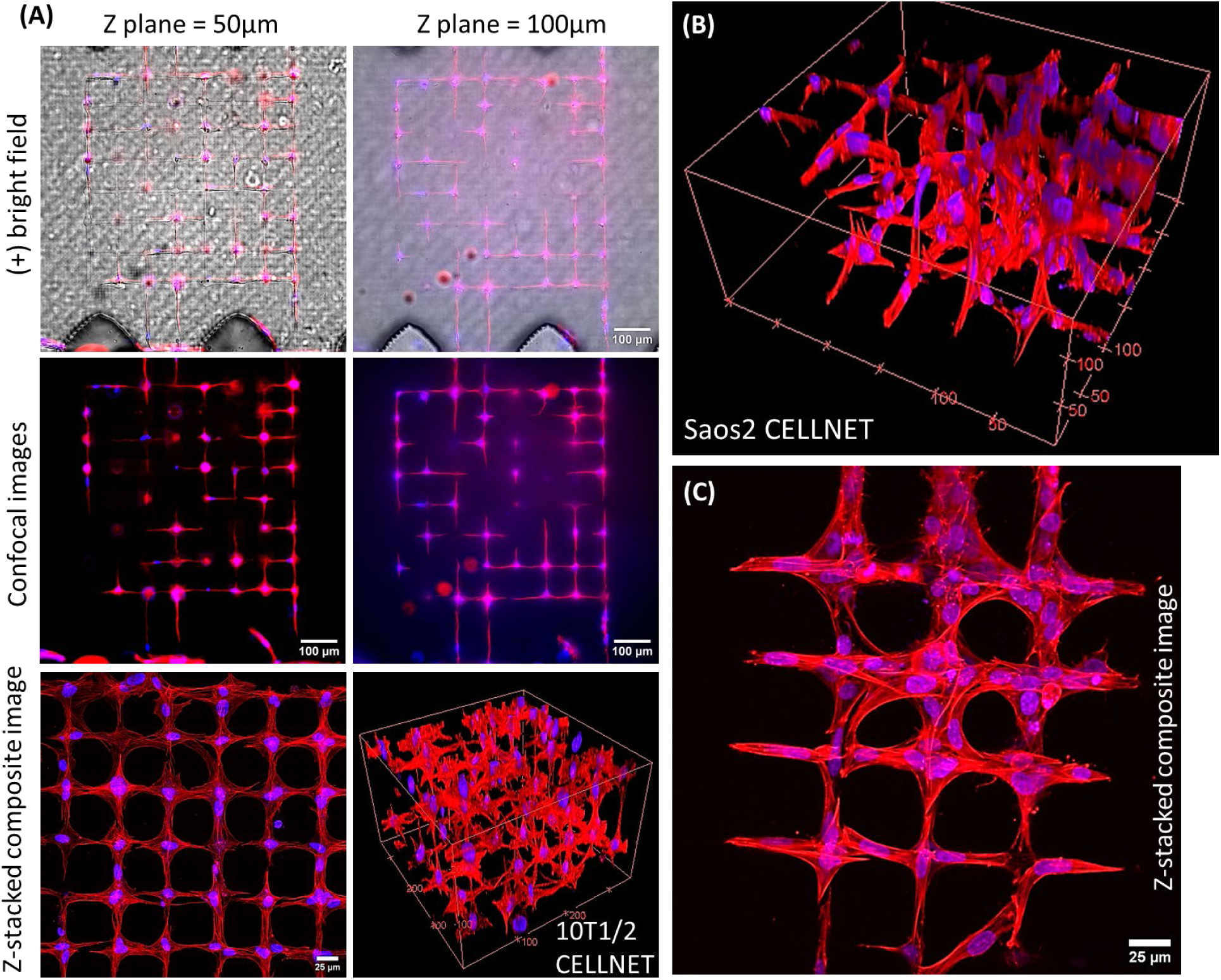
(A) Composite, fluorescence, reconstructed 3D images showing the morphology of 3D (2-layered) CELLNETs using 10T1/2 fibroblast-like cells. (Lumen size ∼8µm) (B-C) 3D reconstructed confocal image and top-view of z-stacked image of Saos2 osteoblast CELLNETs. (Lumen sizes ∼16µm)

**SI-FIG 6.**
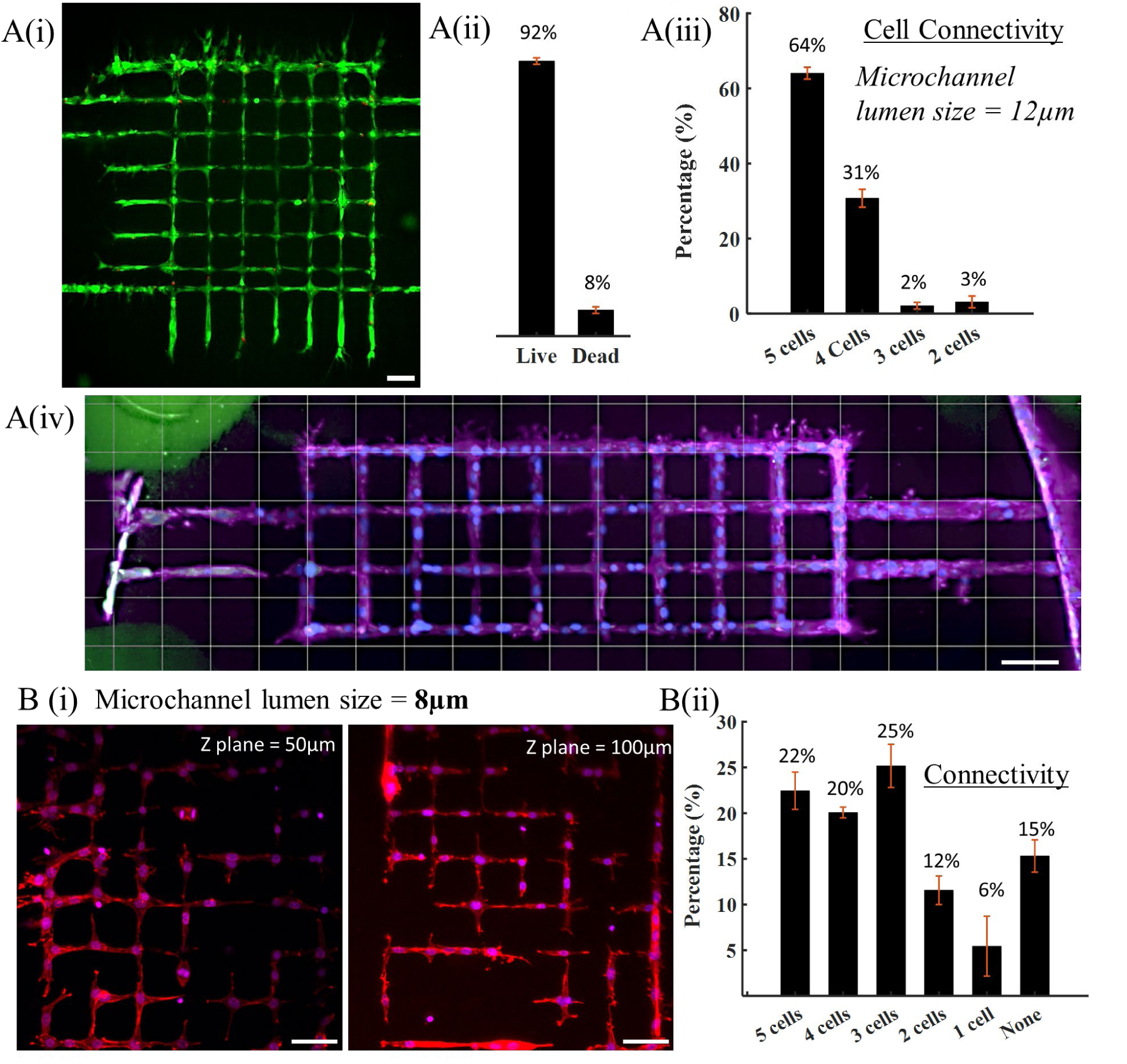
(A) MLO-Y4 CELLNET with lumen size of 8µm. (i) Live/Dead fluroscence image of cell network located ∼50µm inside Type I collagen in Ch#2 of microfluidic devices. (ii) Quantification of live (91.39%) and dead cells (8.61%) from 3 independent devices with ±1.08 (iii) Cell connectivity. Multi cells MLO-Y4 cell circuit Histogram values. Values in x axis are the number of cells connected to each other (5 cells: 64.06% error bar: ±1.92%, 4 cells: 30.73% error bar: ±2.39%, 3 Cells: 2.08% error bar: ±0.90%, 2 cells: 3.13 error bar: ±1.56%) (iv) Stitched images showing MLO-Y4 network spanning the entire width of Ch#2. **(B)** MLO-Y4 CELLNET with lumen size of ∼12µm. (i) Actin Nuclei-stained MLO-Y4 cells at different z planes (ii) Cell Connectivity: Single cell MLO-Y4 cell circuit Histogram values (5 cells: 22.45% error bar: ±2.04%, 4 cells: 20.07% error bar: ±0.59%, 3 cells: 25.17% error bar: ±2.36%, 2Cells: 11.56% error bar: ±1.56%, 1 cell: 5.44 error bar: ±3.28%, none: 15.31 error bar:±1.77). Scale bar A, B: 50μm

**SI-Fig. 7.**
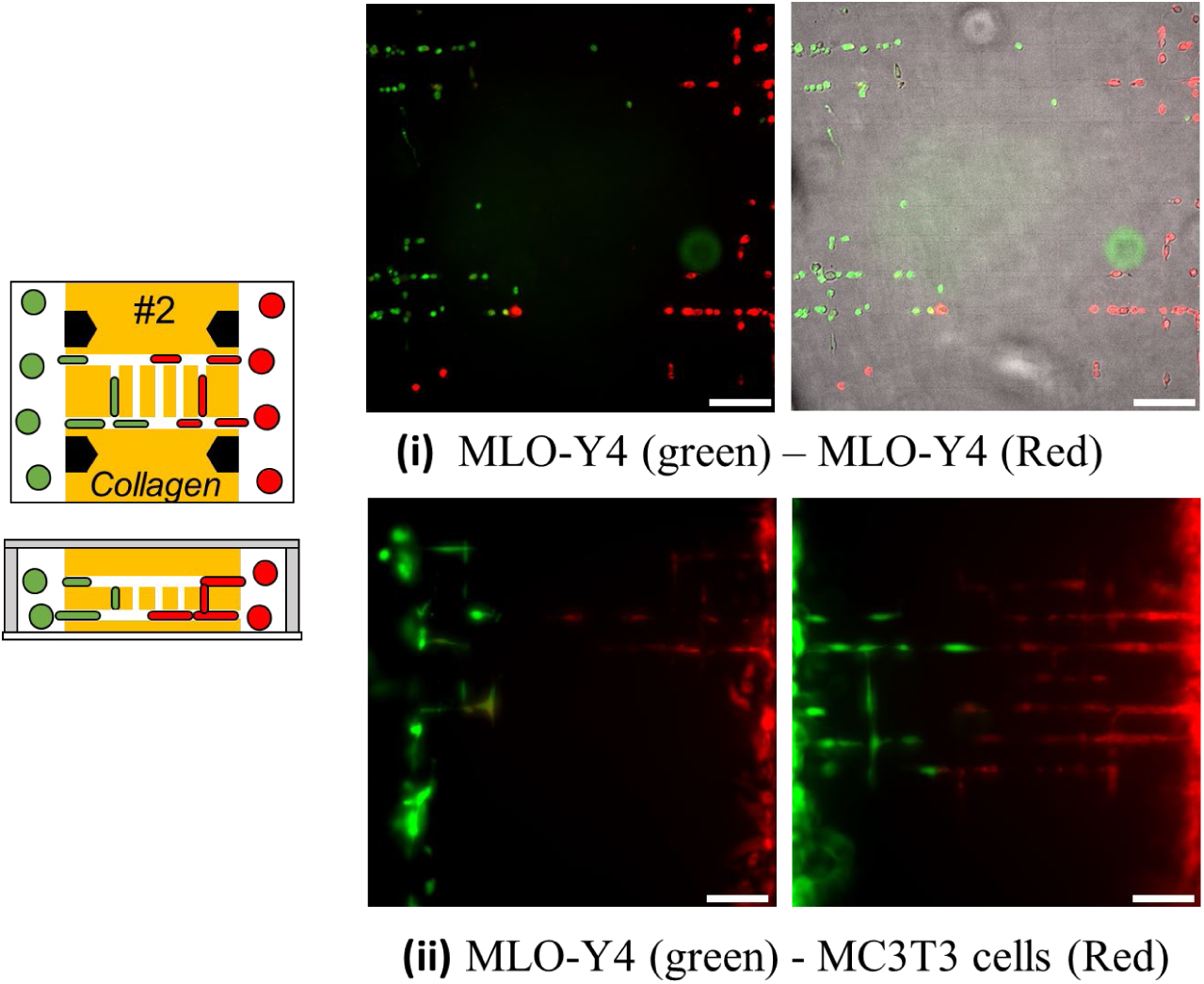
Heterotypic CELLNETs. Scale bar: 100μm.

**SI-Fig. 8.**
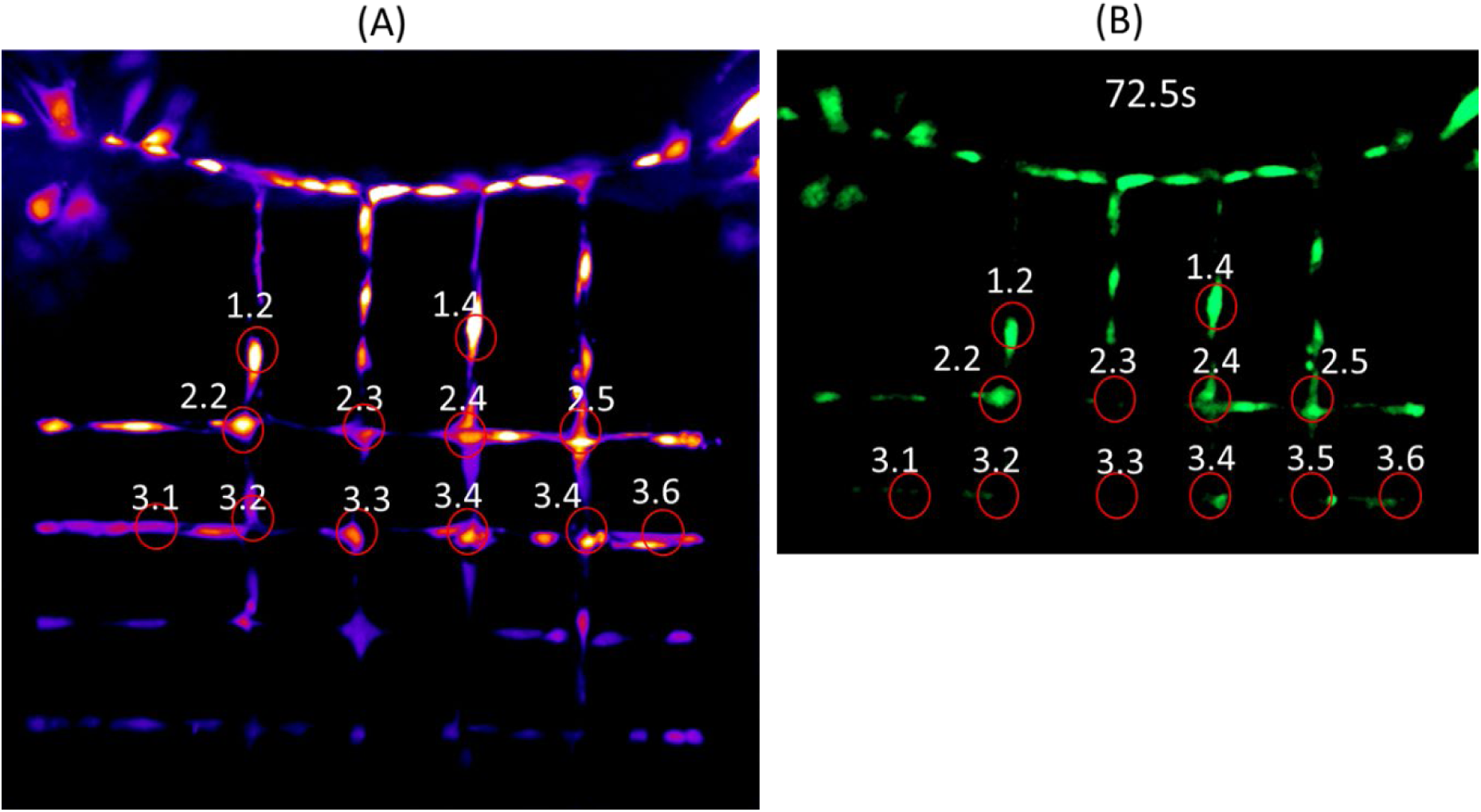
(A) Cumulative heat-map intensity of MLO-Y4 CELLNET during flow-stimuli evoked Calcium signaling experiment. (B) Representative time-lapse fluorescence image showing calcium signaling at specific time-point.

**SI Fig. 9.**
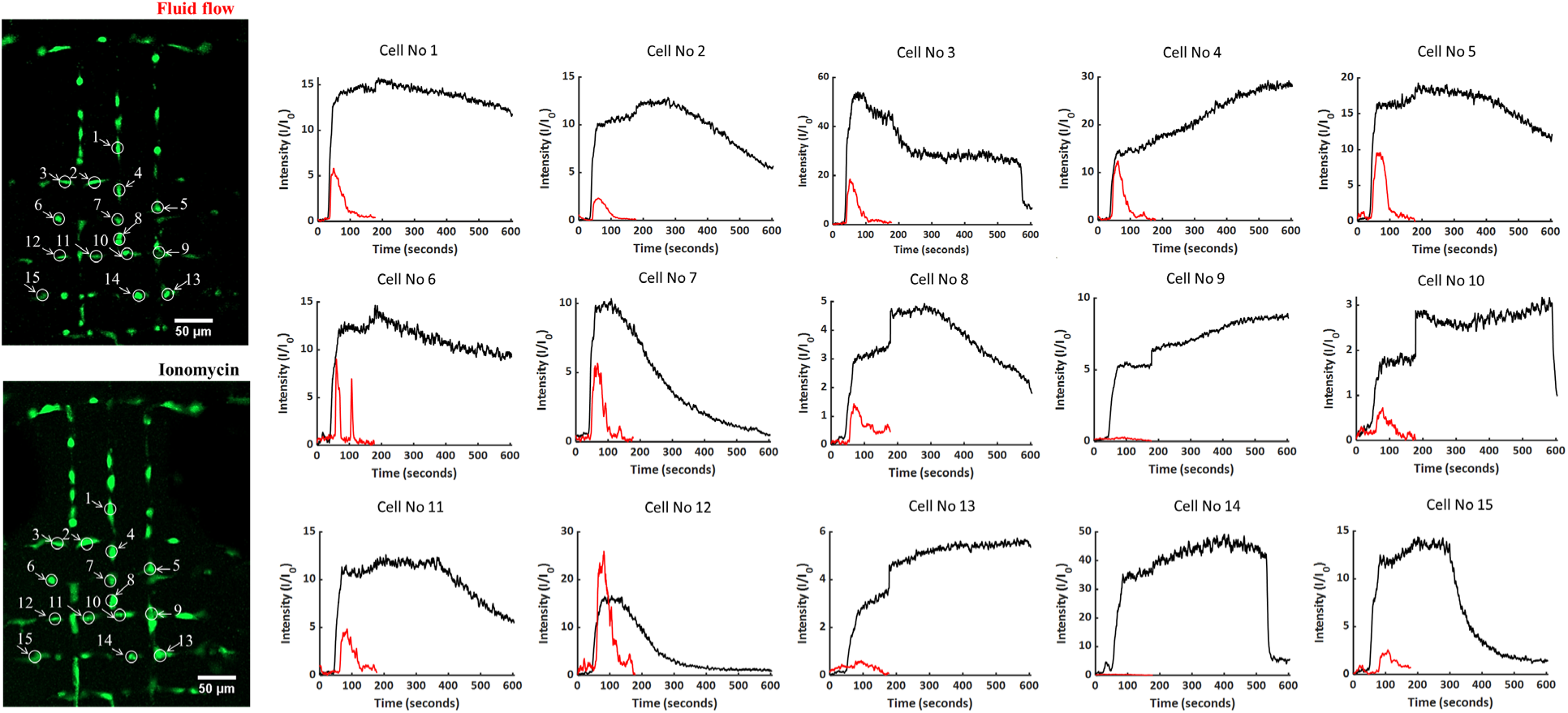
Calcium signaling response of 15 individual osteocytes when subjected to sequential biophysical and biochemical stimuli.

**SI-Fig 10.**
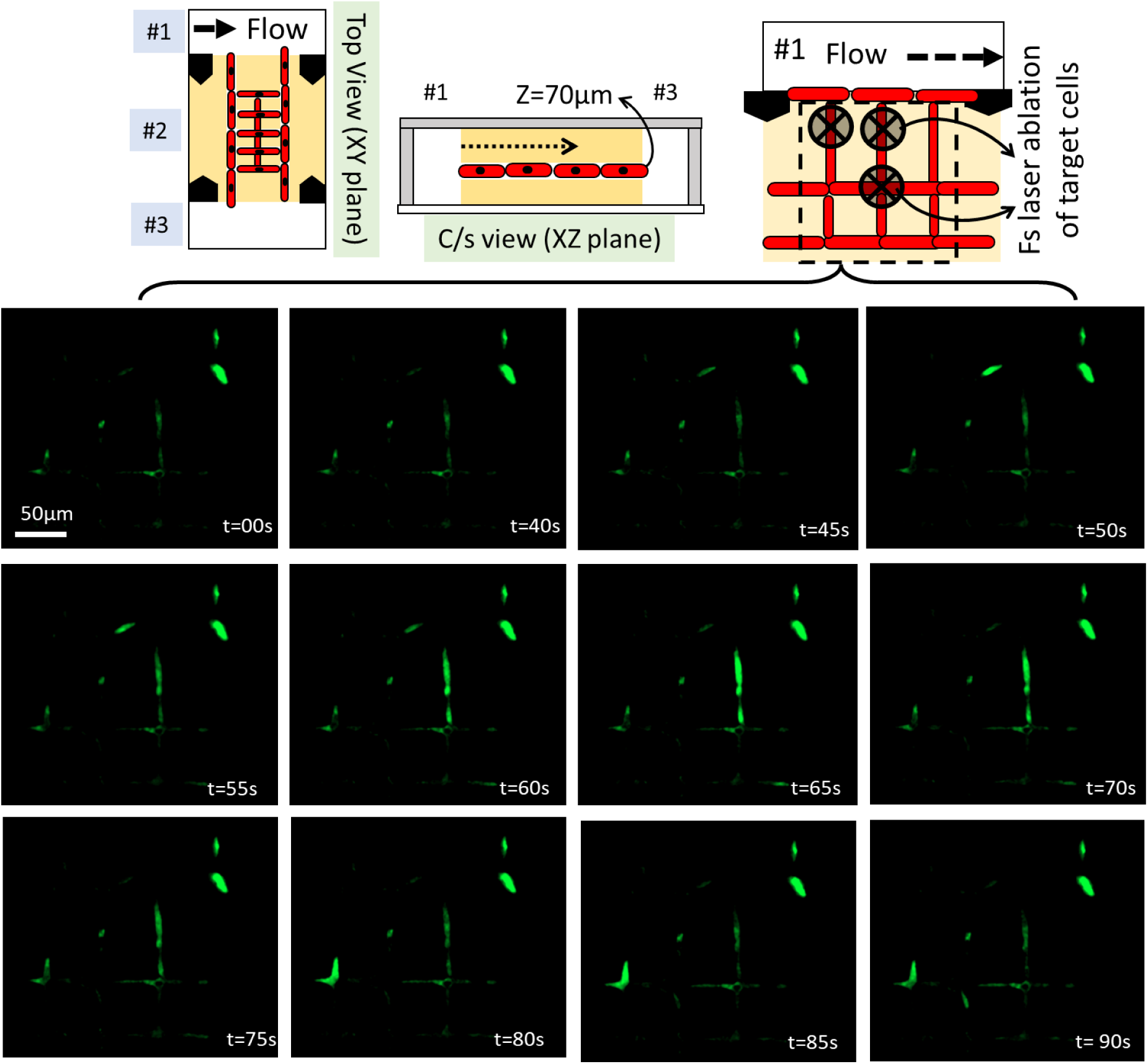
Representative set of real-time fluorescence images show an absence of Ca signal (Fluo-4) in ablated cells forcing signaling through non-ablated connected cells within the network.

**SI-Fig 11.**
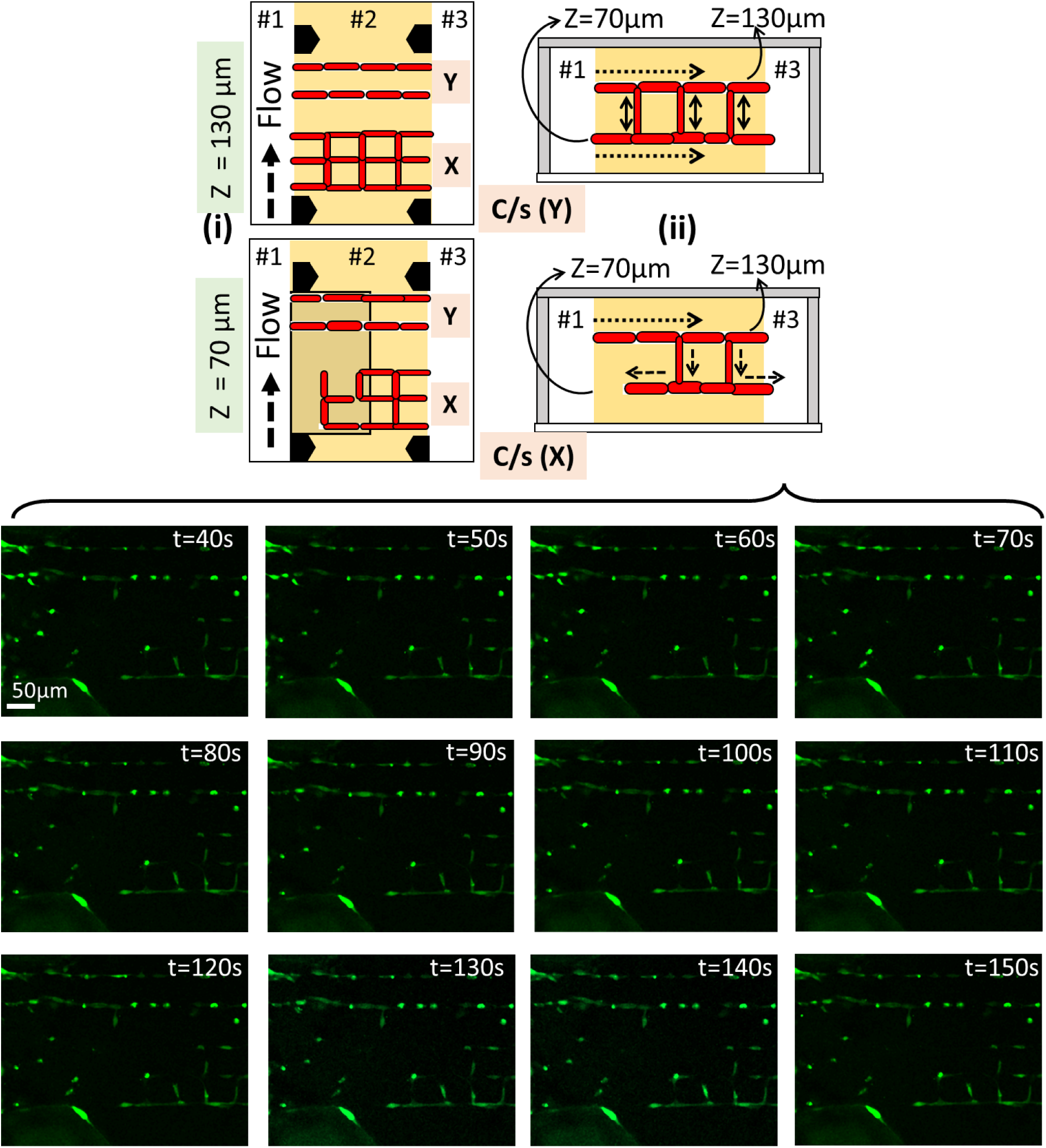
Representative snapshot of real-time fluorescence images showing Ca signaling within the cell network located at z=70µm in a 2-layer osteocyte network with both parallel-line (Y) and square-grid (X) pattens.

**SI-FIG 12.**
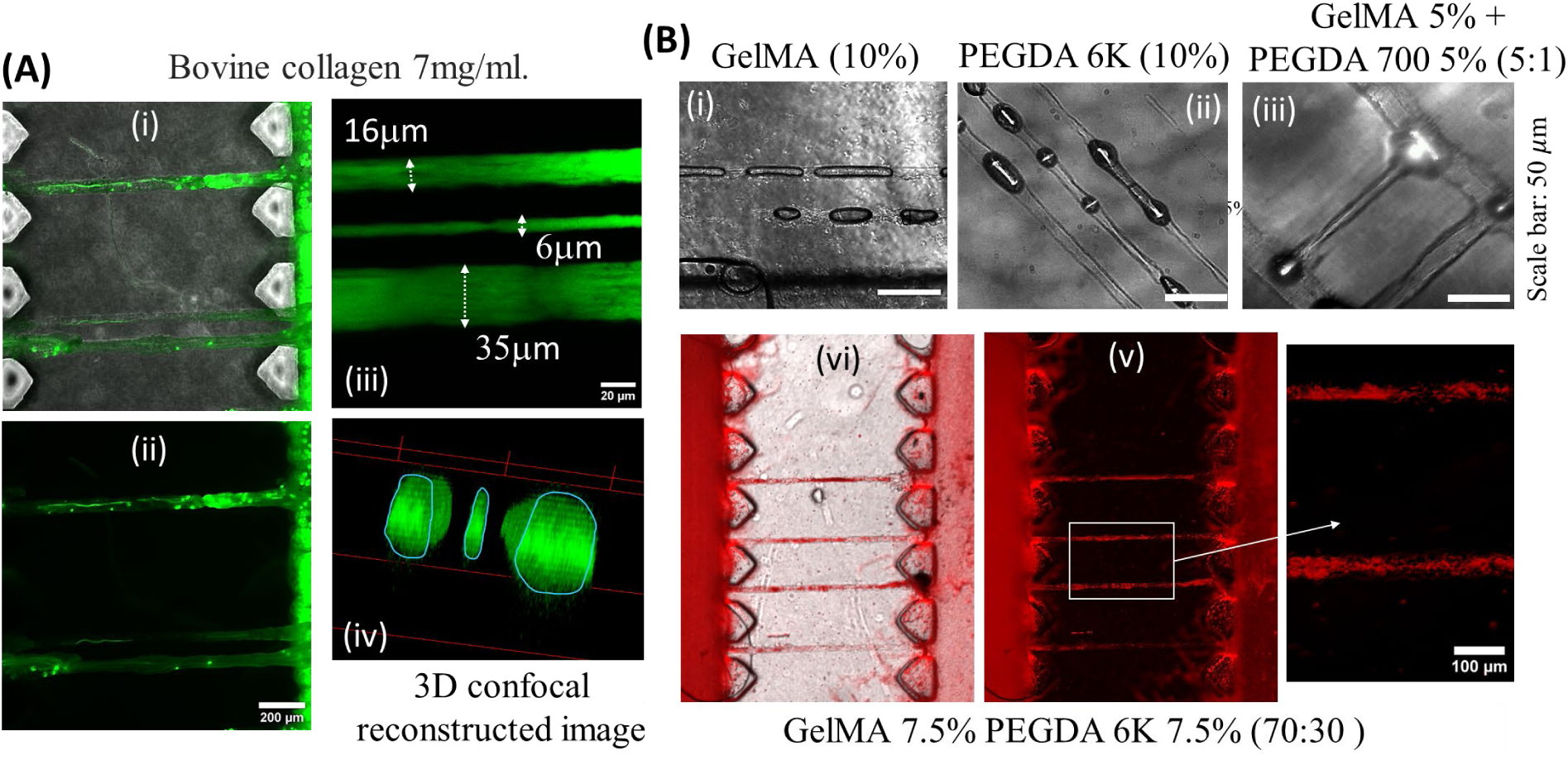
Femtosecond laser ablation was performed in variety of bioinks crosslinked within chamber #2 of PDMS multi-chambered microfluidic devices. (A) Composite (i) and fluorescence (ii) images showing perfusion of green microbeads though ablated channels in bovine collagen; (iii-iv) top and reconstructed isometric views obtained using confocal microscopy showing lumen sizes from 6µm to 35µm. (B) Brightfield, composite and fluorescence images showing perfusable channels within photo-crosslinked hydrogels; (i) Gelatin Methacrylate (GelMA), (ii) Polyethylene glycol diacryate (PEGDA, 6k MW), (iii) a mixture of GelMA-PEGDA, (iv-v) perfusion of red microbead solution through ablated channels in hybrid GelMA-PEGDA hydrogel in Ch#2 of the device.

**SI-Video-1.**
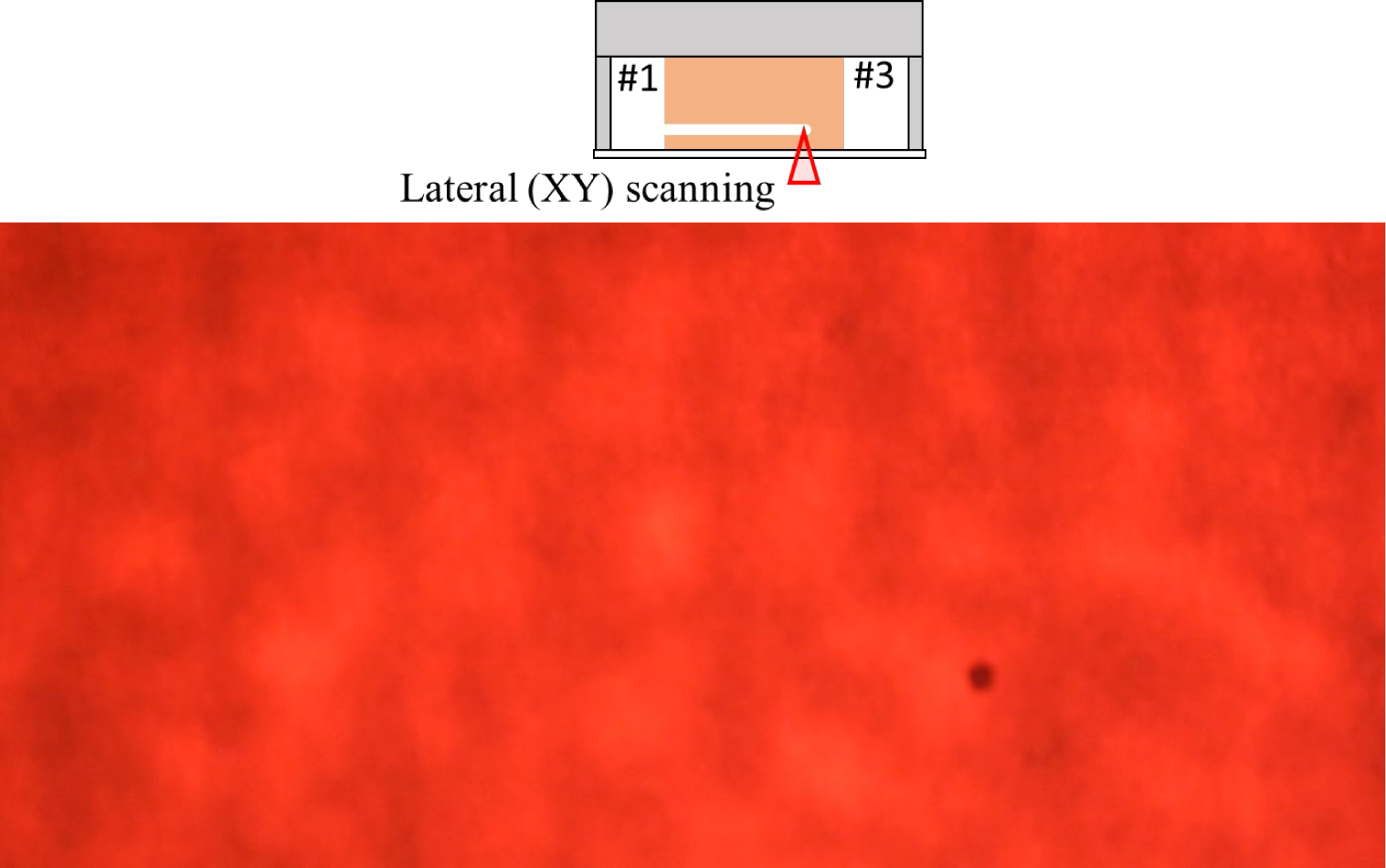
Lateral (XY) scanning in Type I collagen crosslinked within Ch#2 of a multi-chambered PDMS microfluidic device.

**SI-Video-2.**
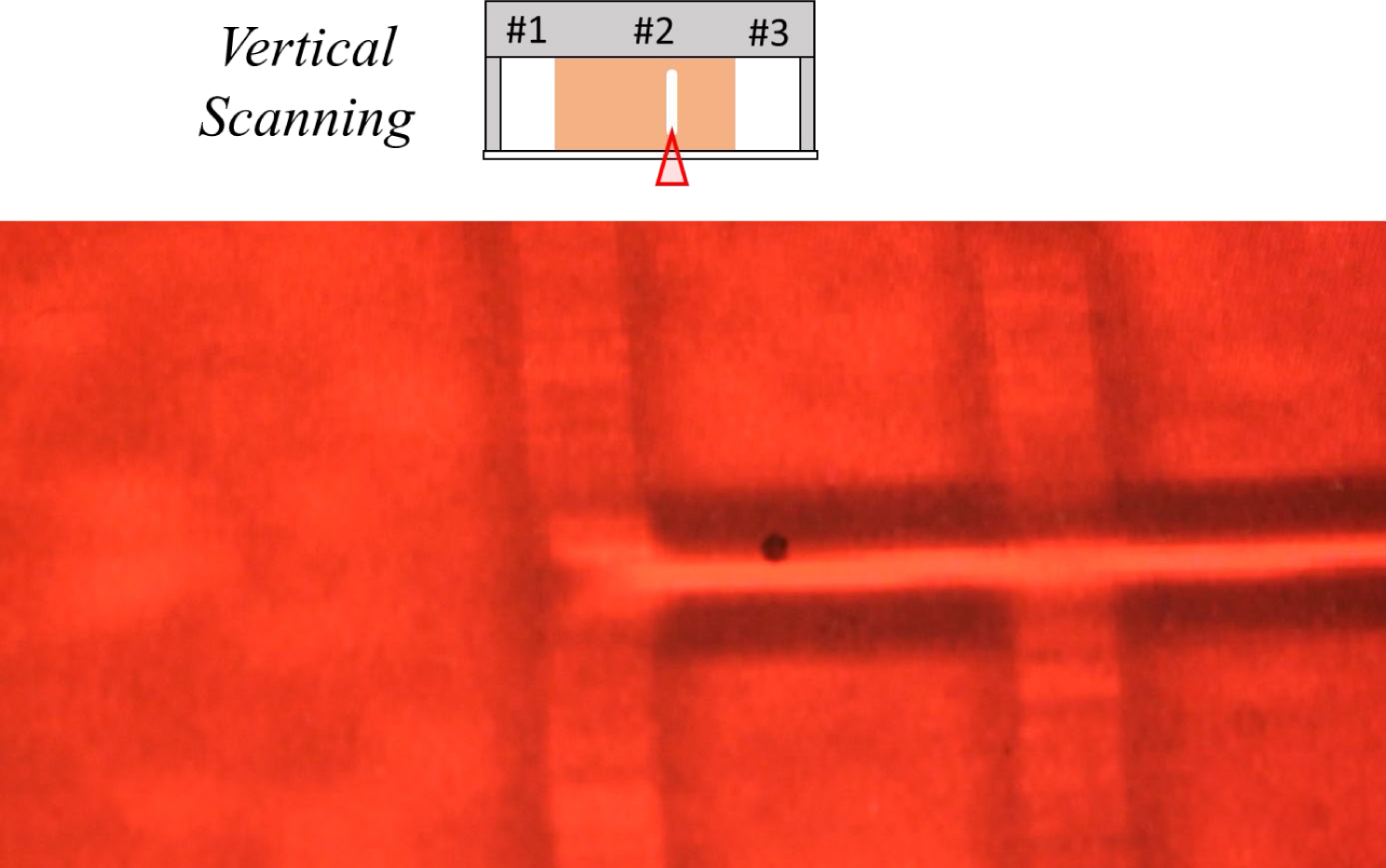
Vertical (Z) scanning in Type I collagen crosslinked within Ch#2 of a multi-chambered PDMS microfluidic device.

